# Endoplasmic reticulum stress inhibition preserves mitochondrial function and cell survival during early onset of isoniazid-induced oxidative stress

**DOI:** 10.1101/2024.08.12.607527

**Authors:** Truong Thi My Nhung, Nguyen Ky Phat, Trinh Tam Anh, Tran Diem Nghi, Nguyen Quang Thu, Ara Lee, Nguyen Tran Nam Tien, Nguyen Ky Anh, Kimoon Kim, Duc Ninh Nguyen, Dong-Hyun Kim, Sang Ki Park, Nguyen Phuoc Long

## Abstract

A comprehensive understanding of isoniazid (INH)-mediated hepatotoxic effects is essential for developing strategies to predict and prevent severe liver toxicity in tuberculosis treatment. Our study utilized multi-omics profiling to investigate the toxic effects of INH, revealing significant involvement of endoplasmic reticulum (ER) stress, mitochondrial impairment, redox imbalance, and altered metabolism. Followed-up mechanistic studies revealed that INH triggered the generation of cytosolic reactive oxygen species (ROS) and the activation of the Nrf2 signaling pathway prior to mitochondrial ROS accumulation. Subsequently, INH disrupted mitochondrial function by impairing respiratory complexes I-IV and caused mitochondrial membrane proton leaks without affecting ATP synthase activity, together leading to mitochondrial depolarization and reduced ATP production. These disturbances enhanced mitochondrial fission and mitophagy. While much attention has been given to mitochondrial dysfunction and oxidative stress in INH-induced hepatotoxicity, our findings highlight the potential of inhibiting ER stress during early INH exposure to mitigate cytosolic and mitochondrial oxidative stress. We further revealed the critical role of Nrf2 signaling in protecting liver cells under INH-induced oxidative stress by maintaining redox homeostasis and enabling metabolic reprogramming via regulating the expression of antioxidant genes and cellular lipid abundance. We also identified other antioxidant pathways (e.g., selenocompound metabolism, HIF-1 signaling pathway, and pentose phosphate pathway) as potential alternative mechanisms besides Nrf2 signaling in response to INH-induced oxidative stress. In conclusion, our research emphasizes the importance of ER stress, redox imbalance, metabolic changes, and mitochondrial dysfunction underlying INH-induced hepatotoxicity.

## Introduction

Tuberculosis (TB) remains one of the leading causes of mortality, accounting for approximately 1.3 million deaths in 2022 [1]. TB treatment is a long-term and complicated procedure, with a standard regimen for drug-susceptible TB requiring at least six months and often associated with various adverse drug reactions [2–4]. Isoniazid (INH), a first-line anti-TB drug, plays a vital role in this regimen [5]. However, severe adverse events, particularly hepatotoxicity, are a significant concern after prolonged use of INH [6]. Therefore, it is crucial to gain more insights into the mechanistic understanding of INH-induced hepatotoxicity.

Several studies highlight the emerging role of mitochondrial dysfunction in the mechanisms underlying INH-induced hepatotoxicity [7]. INH has been shown to cause mitochondrial dysfunction, including impaired mitochondrial complexes, reduced mitochondrial respiration, and diminished energy production [8]. Specifically, INH impairs mitochondrial complexes I, II, and III [9] and mediates the generation of reactive oxygen species (ROS), leading to mitochondrial oxidative stress and apoptosis [10]. Additionally, INH affects mitochondrial biogenesis and dynamics [11]. However, conflicting reports exist, with some studies suggesting that INH does not affect mitochondrial complex I and II activity [8]. While INH alone has been found to reduce whole liver glutathione (GSH) levels and increase lipid peroxidation, it does not alter mitochondrial GSH, oxidized glutathione (GSSG) content, or lipid peroxidation [10], nor does it induce liver cell death [8, 12, 13]. These contradictions raise questions about whether mitochondrial oxidative stress and dysfunction are the primary mechanisms through which INH triggers liver damage.

Nuclear factor erythroid-derived factor 2-related factor 2 (Nrf2) is crucial for cellular defense against oxidative stress and for regulating antioxidant and cytoprotective genes. In the normal cell state, Nrf2 is kept inactive in the cytoplasm by Keap1 [14, 15]. However, under oxidative stress, ROS modify Keap1, Nrf2 moves to the nucleus [16], where it binds to DNA and triggers the production of detoxifying enzymes like glutathione S-transferase (GST), heme oxygenase-1 (Hmox1), and superoxide dismutase (SOD). These enzymes neutralize ROS and repair damaged proteins, reducing oxidative damage and enhancing cell survival. Nrf2 also helps regenerate antioxidants like glutathione. It is reported that Nrf2 can upregulate the expression of SLC7A11, a component of glutamate-cystine antiporter (xCT), which increases cystine uptake, inducing glutathione synthesis [17]. In zebrafish, expression of *Nrf2* and its target genes (*HMOX1*, *NQO1*, *GCLM*, *GCLC*) were found to be upregulated upon INH exposure [18]. However, the role of Nrf2 signaling in INH-induced liver toxicity has yet to be fully elucidated.

Our study provided a more comprehensive analysis of INH-associated hepatotoxicity. Previous research has often examined isolated mechanisms in different biological systems, resulting in fragmented insights. In contrast, our study utilizes multi-omics analysis to offer an integrated view of the metabolic alterations and their association with oxidative stress induced by INH toxicology. We captured the complex interplay between key molecular events, including endoplasmic reticulum (ER) stress, redox imbalance, mitochondrial dysfunction, and cellular metabolic reprogramming. Furthermore, by investigating INH hepatotoxicity in Nrf2-knockout (Nrf2-KO) cells, we highlighted the critical role of Nrf2 signaling and identified alternative protective mechanisms against INH-induced oxidative stress. Our findings present a more complete picture of the molecular mechanisms underlying INH-associated hepatotoxicity and suggest potential targets for both preventive and therapeutic interventions for INH-induced liver injury.

## Material and Methods

### Reagents and Plasmid Construction

Detailed information regarding antibodies, chemicals, and plasmid construction is provided in

### Supplementary Material and Methods. Cell culture and transfection

HepG2, Hep3B, and Huh-7 cell lines were cultured in MEM or DMEM media supplemented with 1% (*v/v*) penicillin-streptomycin (Hyclone, SH40003.01), 1% (*v/v*) sodium pyruvate (Gibco, 11360070), and 10% (*v/v*) FBS (Sigma, 12003C) and maintained in a 5% CO₂ humidified incubator at 37°C. Cells were transfected using Lipofectamine 2000 in OptiMEM2000, following the manufacturer’s protocol, and media were replaced 12 hours post-transfection.

### Generation of the Nrf2 KO HepG2 cell line

Nrf2-KO cells were created using single-guide RNAs (sgRNAs) inserted into the gRNA_Cloning Vector (Addgene, #41824). The sgRNA sequences were GCGACGGAAAGAGTATGAGC and TATTTGACTTCAGTCAGCGA. The efficiency of these sgRNAs was verified by observing GFP-expressing cells using pCAG-EGxxFP (Addgene, #50716), which contained 500 bp of genomic DNA, including the gRNA target sites. Following sgRNAs and Cas9 transfection, cells were subjected to a 4-d selection period using Geneticin, with the dosage determined by an antibiotic kill curve (0.5 mg/mL). Subsequently, cells were plated at single-cell density and allowed to develop for 10-14 d. KO clones were identified and selected through immunoblotting analysis.

### IC20 and IC50 determination

Cells were seeded in 96-well plates at 4×10⁴ cells per well and treated with varying concentrations of Isoniazid (500 µM to 75 mM) for 72 hours, with media refreshed every 24 hours. After treatment, 10 μL of CCK-8 reagent was added to each well, and OD at 460 nm was measured to determine cell viability. IC20 and IC50 values were calculated using dose-response curves for each cell line.

### Cellular and molecular imaging assays

For detailed descriptions of other methods, including ROS detection, immunoblotting, laser scanning confocal microscopy, lipid droplet staining, and immunocytochemistry, please refer to the **Supplementary Material and Methods section**.

### MitoXpress Xtra oxygen consumption assay

Oxygen consumption was monitored using the MitoXpress Xtra Oxygen Consumption Assay, with 1 μM MitoXpress added to each well and sealed with mineral oil. Using a microplate reader (Infinite M200, TECAN), fluorescence was measured at 37°C, with signals recorded for 30 minutes to establish basal respiration, followed by sequential addition of oligomycin (1 μg/mL), FCCP (2 μM), and rotenone/antimycin A (1 μM each), recording signals for 30 minutes after each addition. Measurements were normalized to 1000 cells and used to determine basal, ATP-linked, maximal, and non-mitochondrial respiration rates by comparing oxygen consumption rate (OCR) before and after each treatment.

### RNA extraction, cDNA synthesis and RT-qPCR

Cell pellets were subjected to RNA isolation using TRIzol reagent (Virginia Tech Bio-Technology, #TS200-001). cDNA was synthesized using Invitrogen Superscript IV Reverse Transcriptase (Invitrogen, #18090050) and oligo-dT primers (Invitrogen, #18418020). qPCR was conducted with FastStart SYBR Green Master Mix (Roche, #04913850001) on a StepOnePlus™ Real-Time PCR System (Applied Biosystems, #15360337), using GAPDH for normalization via the2^-ΔΔCt^ method. The following primers were used for housekeeping gene and Nrf2 target genes: *GAPDH*, CTCCTGCACCACCAACTGCT and GGGCCATCCACAGTCTTCTG; *GCLC*, TCCAGGTGACATTCCAAGCC and GAAATCACTCCCCAGCGACA; *HMOX1*, AGTCTTCGCCCCTGTCTACT and GCTGGTGTGTAGGGGATGAC; *NQO1*, GCACTGATCGTACTGGCTCA and CCACCACCTCCCATCCTTTC; *SLC7A11*, TTTTCTGAGCGGCTACTGGG and CAGCTGGTAGAGGAGTGTGC.

### High-throughput cellular omics experiments

Cellular extraction of metabolites, lipids, and proteins for metabolomics, lipidomics, and proteomics data acquisition, as well as data processing and normalization, are described in detail in the **Supplementary Material and Methods**.

### Statistical analysis

Quantitative data are shown as mean ± standard deviation (SD) unless specified differently. Sample sizes differed according to the experimental conditions. Statistical tests were carried out using GraphPad Prism 10.2.3. Two-sided Student’s t-test was used to compare the means between the two groups. To compare means across more than two groups, one-way or two-way ANOVA with Tukey’s post hoc test was applied. A significance level of 0.05 was used.

The log-transformed median-normalized omics data were subject to univariate analyses. The *limma* method [19] was used for single-factor differential analysis to identify differential proteins, while the unpaired two-sided t-test was performed to identify differential metabolites and lipids. Regarding two-factor differential analysis, ExpressAnalyst [20] (for proteomics data) and MetaboAnalyst 6.0 [21] (for metabolomics and lipidomics data), which implemented the limma functionality, were employed. The false discovery rate cut-off of 0.05 was applied to consider the significant differential metabolites/lipids/proteins.

We conducted functional analysis to gain biological insights into molecular pathways from significantly differential proteins, metabolites, or lipids. Detailed information regarding functional analysis can be retrieved from **Supplementary Material and Methods.**

## Results

### INH exposure induces ROS and apoptosis in HepG2, Hep3B, and Huh-7 cells

To explore the cytotoxicity of INH in hepatocytes, we assessed cell viability in three different liver cell lines (HepG2, Hep3B, Huh-7) at various concentrations of INH ranging from 0.5 to 75 mM (Fig. 1A). Using these genetically and functionally diverse cell lines ensures that our findings on INH’s hepatotoxicity are robust and generalizable, confirming that the observed cytotoxic patterns are not restricted to the specific characteristics of a single cell type. After a 72-h incubation, the IC20 and IC50 values of INH for each cell line were determined, representing 20% and 50% reductions in cell viability compared to the control group. IC20 and IC50 were respectively 17 and 32.5 mM for HepG2 WT cells, 15 and 60 mM for Hep3B cells, and 25 and 45 mM for Huh-7 cells. We further examined how IC20 and IC50 doses of INH affected the expression of apoptosis-related proteins, including Bax, Bcl-2, cleaved Caspase-9, and cleaved Caspase-3. Using immunoblotting analysis, we observed that increasing concentrations of INH for 72 h elevated the levels of Bax, cleaved Caspase-9, and cleaved Caspase-3, accompanied by a corresponding decrease in Bcl-2 levels in all three hepatic cell lines (Fig. 1B). These results indicate that INH induces apoptosis in these liver cell lines in a dose-dependent manner.

**Figure 1.**
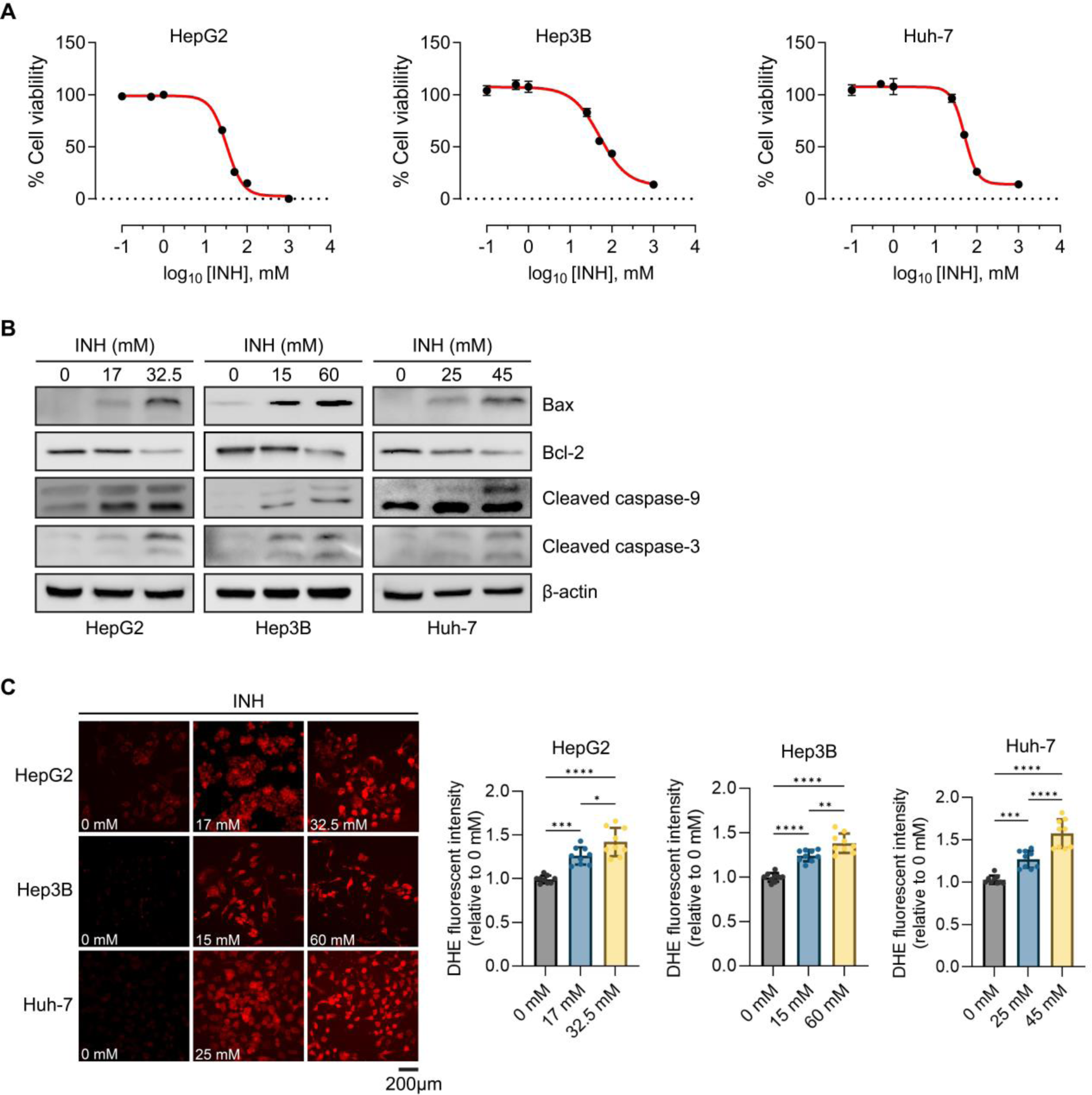
Isoniazid (INH) impaired cell viability and induced apoptosis, associated with elevated cellular ROS levels. (A) Dose-response curve of INH in HepG2, Hep3B, and Huh-7 cells for 72 h. (B) INH exposure influenced the expression of apoptosis-related proteins.

Representative immunoblots of Bax, Bcl-2, cleaved Capase-9, and cleaved Caspase-3 proteins after 72-h INH exposure with concentrations that inhibit 20% and 50% cell growth in three cell lines. β-actin served as protein loading control. (C) Representative immunofluorescence images of DHE staining (left) and quantitative data (right) after INH treatment for 72h in the indicated cells. Data are presented as means ± SD from three independent experiments. Statistical differences between groups were determined by one-way ANOVA with Tukey post-tests for multiple comparisons, ****P-value < 0.0001.

It was reported that INH-induced apoptosis is related to oxidative stress [22, 23], so we utilized dihydroethidium (DHE) staining to investigate the impact of INH on ROS production in these three liver cell lines treated with varying concentrations of INH for 72 h. As shown in Fig. 1C, INH-treated samples in all cell lines exhibited significantly increased ROS production compared to the control group. Altogether, the findings from apoptosis-related protein analysis and DHE staining suggest that oxidative stress is associated with the cytotoxic effects triggered by INH.

### Multi-omics reveals a comprehensive landscape regarding alterations of multiple cellular processes under INH exposure

INH treatments for 72 h significantly altered the proteome of HepG2 WT cells compared to controls (Fig. 2A). Specifically, 1376 and 1855 differential proteins were identified in the 17 mM and 32.5 mM INH treatments versus control comparisons, respectively (Supplementary Fig. S1A-B). Kyoto Encyclopedia of Genes and Genomes (KEGG) pathway enrichment analysis revealed that 32.5 mM INH treatment affected ER unfolded protein response (“Protein processing in endoplasmic reticulum”), redox homeostasis (“Cysteine and methionine metabolism,” “Glutathione metabolism,” “Ferroptosis”), mitochondrial metabolic pathways (“TCA cycle,” “Pyruvate metabolism,” “Oxidative phosphorylation,” and “Fatty acid metabolism”), and ROS-related pathways (“Chemical carcinogenesis - reactive oxygen species”) (Fig 2. B-E, Supplementary Table S1). The 17 mM INH-treated group followed a similar pattern of molecular alterations but to a lesser extent (Fig 2. B-E). The differential proteins were also enriched in Gene Ontology Cellular Component (GO:CC) terms related to ER and mitochondrial complexes (Supplementary Table S2). The abundance of multiple proteins related to ER stress response, including ERO1A, HSPA5, HSP90B1, PDIA3, PDIA4, CANX, CALR, DNAJB11, ERLEC1, HYOU1, and ERP29, were increased (Fig. 2B, Supplementary Fig. S2A), suggesting the involvement of the unfolded protein response (UPR).

**Fig. 2.**
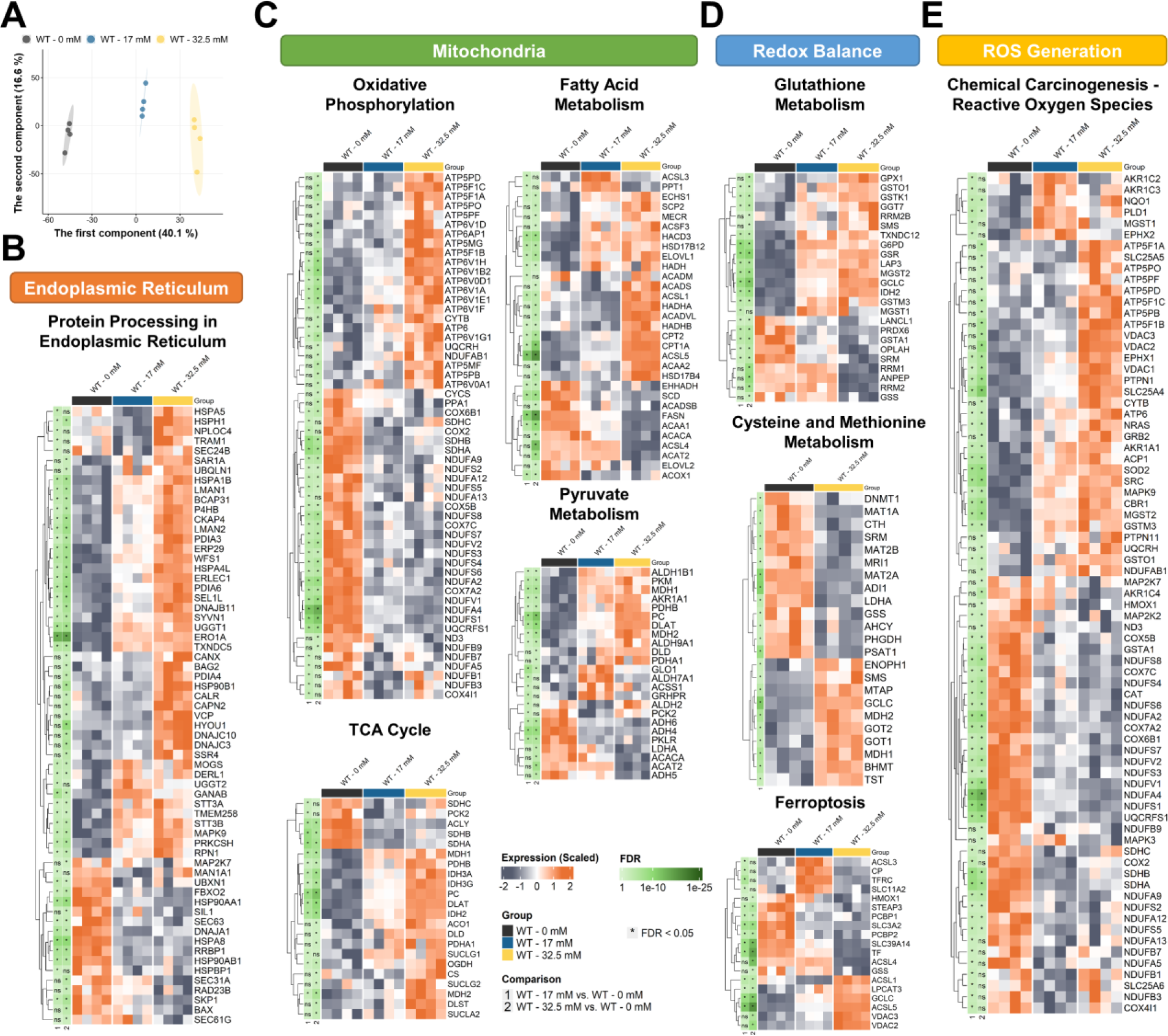
Proteomics analysis suggested that INH treatment induced endoplasmic reticulum stress, mitochondrial dysfunction, oxidative stress, altered redox state, and perturbations of cellular metabolism in HepG2 cells. (A) Principal component analysis scores plot of proteome profiles of INH-treated and untreated WT HepG2 cells. (B-E) Heatmaps of significantly differential proteins between INH-treated and untreated WT HepG2 cells enriched into significant KEGG pathways related to endoplasmic reticulum (B), mitochondria (C), redox balance (D), and reactive oxygen species generation (E). Abbreviation: WT, wild-type; FDR, false discovery rate; KEGG, Kyoto Encyclopedia of Genes and Genomes.

We also observed significant enrichment of mitochondrial protein complexes in both oxidative phosphorylation and ROS-related pathways (Fig. 2C and E, Supplementary Fig. S2B). In particular, the decreased abundance of numerous subunits of complex I (NDUFA2, NDUFA4, NDUFA9, NDUFS1, NDUFS2, NDUFS3, NDUFS4, NDUFV1, among others), II (SDHA, SDHB, SDHC), III (UQCRFS1), IV (COX6B1, COX7A2, COX7C) suggests impaired mitochondrial electron transport chain function, which is essential for ROS induction. Conversely, the increased expression levels of complex V subunits (ATP5F1A, ATP5F1B, ATP5F1C, ATP6V1B2, ATP6V1E1, among others), TCA cycle enzymes (MDH1, MDH2, PDHB, IDH3A, IDH3G, IDH2, PC, ACO1, DLAT, DLD, OGDH, and SUCLG1), enzymes involved in the conversion of pyruvate to acetyl-CoA (PDHB, PDHA1, and DLAT), β-oxidation enzymes (ACSL1, ACSL5, CPT1A, CPT2, HADHA, HADHB, ACADS, and ACADVL) were observed. Remarkably, these molecules are all involved in ATP production and maintenance of energy homeostasis. Moreover, we found higher abundance of proteins for glutathione synthesis and antioxidant defense (GPX1, GSTK1, GSTO1, G6PD, GSR, and GCLC), ferroptosis (ACSL1 and ACSL5), detoxifying enzymes in response to oxidative stress (SOD2, AKR1C2, AKR1C3, NQO1, MGST1, MGST2, and GSTM3). These findings suggest the alterations in glutathione production, lipid peroxidation, and redox imbalance due to oxidative stress caused by INH treatment over 72 h (Fig. 2D-E). Collectively, proteomic data suggest the changes in ER stress response, mitochondrial complex I-IV dysfunction, along with metabolic responses to stress conditions induced by INH.

Since metabolic alterations were found at the proteome level, we further performed high-throughput metabolic profiling to examine the consequences of INH treatment in cellular metabolism via metabolomics and lipidomics. In the principal component analysis scores plot, the metabolome of the HepG2 WT cells receiving INH treatment showed significant separation from the control group (Fig. 3A). Compared to the control, the 17 mM INH-treated group had 20 significantly differential metabolites, while 32.5 mM INH-treated group had 32 significantly differential metabolites (Supplementary Fig. S3A-B). Despite the increased abundance of TCA cycle enzymes and proteins associated with oxidative stress response, metabolites related to the TCA cycle (e.g., citrate, isocitrate, malate, and beta-nicotinamide adenine dinucleotide) and glutathione metabolism (e.g., GSH, GSSG, glutamine) were found to be significantly decreased in 32.5 mM INH-treated group (Fig. 3B). The 17 mM INH-treated group showed a similar pattern with a lesser extent in significant metabolites (Fig. 3B). In addition, results of pathway analysis showed that 17 mM INH treatment was associated with a KEGG pathway related to redox balance (“Glutathione metabolism”), and 32.5 mM INH treatment was associated with a pathway related to mitochondria [“Citrate cycle (TCA cycle)”] (Supplementary Table S3). These findings suggested a disruption in the TCA cycle and an altered redox state due to the toxic effects of INH treatment for 72 h, even with the enhanced expression of TCA enzymes and antioxidant proteins, which might be the compensatory mechanisms for maintaining cellular homeostasis. Besides, in both 17 mM and 32.5 mM INH-treated groups, the abundance of carnitine and acylcarnitine with shorter chain [CAR(2:0)] were decreased, whereas the abundance of acylcarnitines with longer chain [CAR(16:0) and CAR(14:0)] was increased compared to controls (Fig. 3B).

**Fig. 3.**
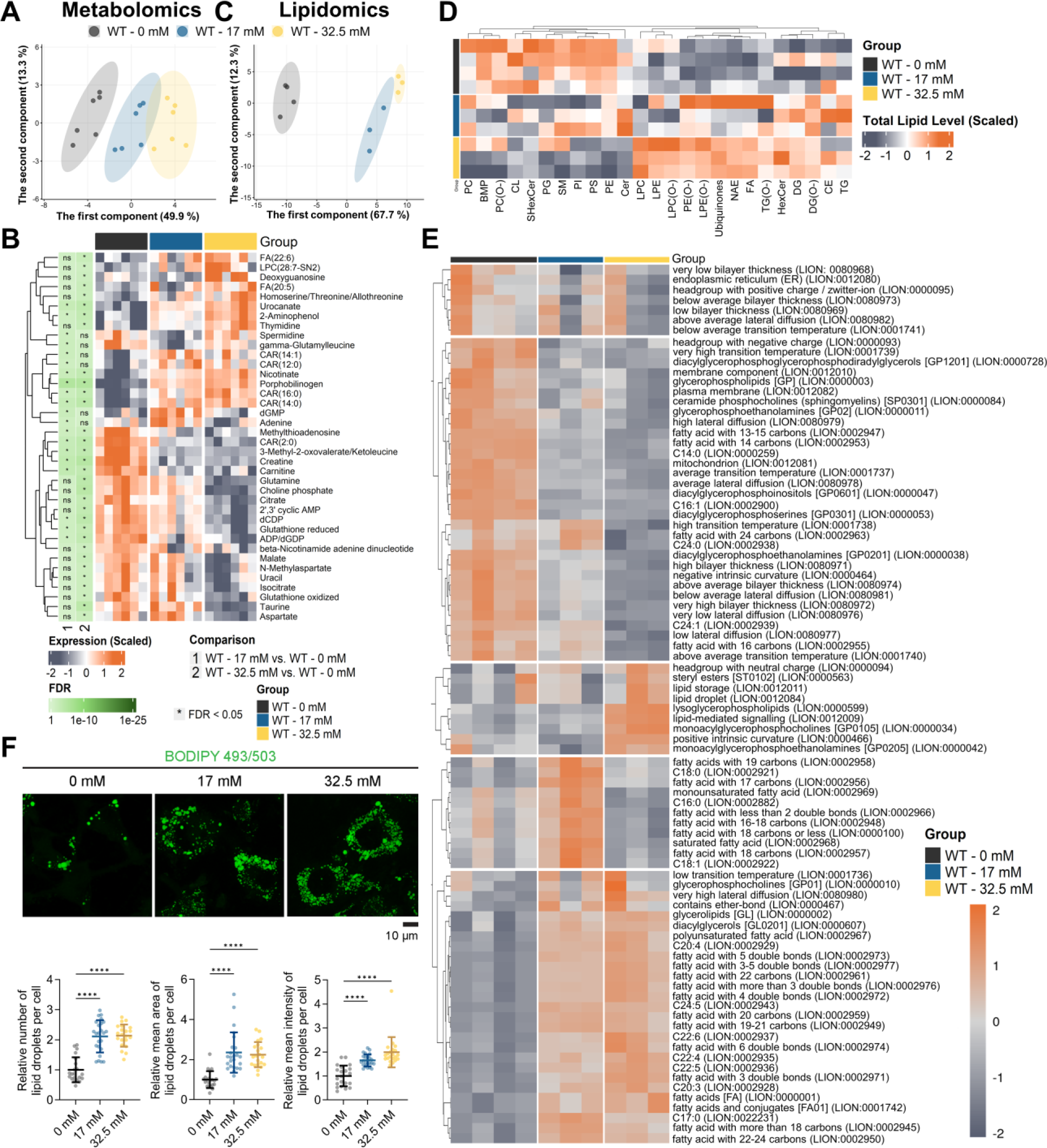
Metabolomics and lipidomics analyses suggested cellular perturbations related to mitochondrial dysfunction, oxidative stress, and lipid droplet formation in HepG2 cells under INH exposure. (A) PCA scores plot of metabolome profiles of INH-treated and untreated WT HepG2 cells. (B) Heatmap of significantly differential metabolites between INH-treated WT HepG2 cells and controls. (C) PCA scores plot of lipidome profiles of INH-treated and untreated WT HepG2 cells. (D) Heatmap presents differences in total lipid subclass levels between INH-treated WT HepG2 cells and controls. (E) PCA heatmap of lipid ontology enrichment analysis of significantly differential lipids between INH-treated WT HepG2 cells and controls. (F) Representative images of INH-treated HepG2 cells stained with BODIPY 409/503 (left) and quantification of lipid droplet (LD) accumulation in cells (right). Both concentrations of INH increased LD number, size (area), and intensity in HepG2 cells after 72 h of treatment. Data are shown as means ± SD, analyzed from more than 100 cells per group. Statistical differences between groups were determined by one-way ANOVA with Tukey post-tests for multiple comparisons; ****P < 0.0001. Abbreviation: PCA, principal component analysis, WT, wild-type; FDR, false discovery rate; PC, Phosphatidylcholine; BMP, Bismonoacylglycerophosphate; PC(O-), Ether-linked phosphatidylcholine; CL, Cardiolipin; SHexCer, Sulfatide; PG, Phosphatidylglycerol; SM, Sphingomyelin; PI, Phosphatidylinositol; PS, Phosphatidylserine; PE, Phosphatidylethanolamine; Cer, Ceramide; LPC, Lysophophatidylcholine; LPE, Lysophosphatidylethanolamine; LPC(O-), Ether-linked lysophosphatidylcholine; PE(O-), Ether-linked phosphatidylethanolamine; LPE(O-), Ether-linked lysophosphatidylethanolamine; NAE, N-acyl ethanolamines; FA, Fatty acid; TG(O-), Ether-linked triacylglycerol; HexCer, Hexosylceramide; DG, Diacylglycerol; DG(O-), Ether-linked diacylglycerol; CE, Cholesteryl ester; TG, Triacylglycerol.

Regarding the lipidomics analysis, 17 mM and 32.5 mM INH-treated groups showed a distinct lipid profile compared to the control group (Fig. 3C) with 281 and 358 significantly differential lipids, respectively (Supplementary Fig. S3C-D). The total lipid levels of cholesteryl ester (CE), diacylglycerol (DG), triacylglycerols (TGs), and fatty acids (FAs) were generally higher in the treatment group, where the 32.5 mM clearly showed a more considerable extent of significant lipid species (Fig. 3D, Supplementary Fig. S4). On the contrary, total lipid levels of sphingomyelin (SM), phosphatidylethanolamines (PEs), phosphatidylserines (PSs), phosphatidylinositols (PIs), phosphatidylcholines (PCs), cardiolipins (CLs), and ether-linked phosphatidylcholines [PC(O-)] were lower in the treatment groups and also exhibited a dose-dependent pattern (Fig. 3D, Supplementary Fig. S4). The enrichment analysis indicated a dose-dependent reduction in the abundance of lipid species associated with ER, mitochondria, and membrane components in INH-treated groups (Fig. 3E). Moreover, the INH-treated groups also displayed an upregulation of polyunsaturated fatty acids (PUFAs) and lipid species related to the formation of lipid droplets (Fig. 3E). The elevation in the abundance of PUFAs, FAs, and acylcarnitine, combined with the reduction of carnitine level, indicated the disruption of the β-oxidation process. Additionally, these observations suggested that the carnitine shuttle pathway, which plays a key role in the carnitine and long-chain fatty acids transport through mitochondrial membranes for β-oxidation, was impaired. The increased expression levels of proteins involved in converting pyruvate to acetyl-CoA might compensate for the shortage of acetyl-CoA due to disrupted β-oxidation. In agreement with our lipidomics data, we observed an increase in the number, size, and intensity of lipid droplets in INH-treated cells after 72 h using BODIPY staining. However, no significant difference was noted between the two concentrations (Fig. 3F).

Overall, our results from multi-omics data analysis indicated that exposing HepG2 WT cells to INH was associated with 4 critical molecular events: ER stress, mitochondrial dysfunction, disrupted redox imbalance and oxidative stress, and significantly altered metabolism.

### INH induces mitochondrial dysfunction and stress responses in HepG2 WT cells

We first explored the effects of 72 h of INH treatment on mitochondria by quantifying mitochondrial morphology in HepG2 WT cells (Fig. 4A-E). INH significantly decreased the mitochondrial content and area (Fig. 4B-C), reduced the aspect ratio (Fig. 4D), and increased the number of individual mitochondria (Fig. 4E). These data suggest that INH enhanced mitochondrial fission and mitophagy.

**Fig. 4.**
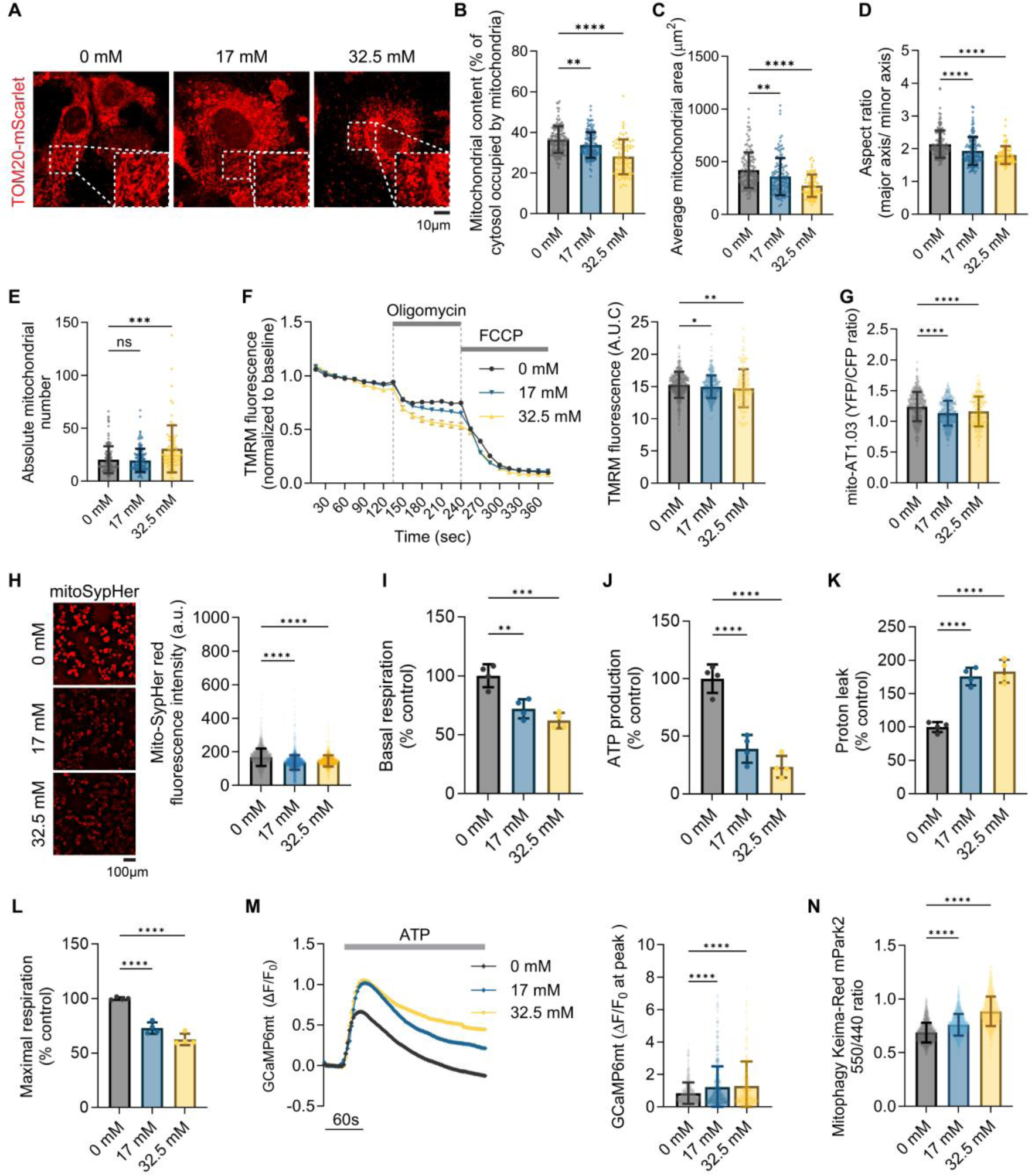
INH exposure resulted in mitochondrial dysfunction after 72 h. (A-E) Mitochondrial morphology analysis in HepG2 cells treated with INH for 72 h. (A) Representative images of mitochondrial networks using TOM20-mScarlet in HepG2 cells treated with 0 mM, 17 mM, and 32.5 mM INH. Magnified views of the boxed regions in the main images (bottom left) highlight changes in mitochondrial structure. Scale bar: 10 µm. (B) Quantification of mitochondrial content is expressed as the percentage of cytosol occupied by mitochondria. (C) Measurement of average mitochondrial area. (D) Aspect ratio analysis, representing the ratio of major to minor axes of the mitochondria. (E) The absolute number of mitochondria per cell. (F) TMRM fluorescence intensity over time was normalized to baseline, indicating mitochondrial membrane potential in the presence of oligomycin and FCCP. (G) Mito-AT1.03 YFP/CFP fluorescence ratio, indicating ATP levels in mitochondria. (H) Representative images and quantification of mitoSypHer-red fluorescence intensity, showing mitochondrial pH. Scale bar: 2 mm. (I-L) Mitochondrial oxygen consumption rate determined using mitoXpress of HepG2 after 72-h INH treatment including (I) basal respiration rate, (J) ATP production rate, (K) proton leak, and (L) maximal respiration compacity. (M) Monitoring mitochondrial Ca^2+^ uptake in INH-treated HepG2 cells using GCaMP6mt sensor before and after ATP addition (left). Data for the maximum peak of GCaMP6mt signals are shown (right). (N) Quantification of mitoKeima-Red using fluorescence intensity ratio (550/440) in WT HepG2 cells after 72 h of INH exposure. Data are presented as means ± SD, analyzed from more than 100 cells per group. Statistical differences between groups were determined using one-way ANOVA with Tukey’s post-test for multiple comparisons; n.s., not significant, *P-value <0.05, **P-value < 0.01, ***P-value < 0.001, ****P-value < 0.0001.

We monitored mitochondrial membrane potential (Δψm) using TMRM fluorescence (Fig. 4F). INH exposure for 72 h caused immediate depolarization of Δψm upon treatment with the ATP synthase inhibitor oligomycin and a sharp drop with the protonophore FCCP. These results suggest that INH impairs mitochondrial respiration, and to maintain Δψm, mitochondria rely primarily on ATP hydrolysis by complex V. We further found the depletion of mitochondrial ATP production under 72-hour INH treatment using a mitochondria-targeted ATP probe (mitoAT1.03) (Fig. 4G). In addition, mitochondrial acidification under INH exposure was indicated by the reduction in mito-SypHer red signals (Fig. 4H), likely due to increased proton accumulation within the matrix. To determine the impact of INH on mitochondrial function, we measured mitochondrial oxygen consumption rates (OCRs) using mitoXpress at 72 h after treatment. Fig. 4I-L shows that INH-treated groups exhibited lower basal respiration rates, reduced ATP production, and decreased maximal respiratory capacity despite increased proton leakage. These findings suggest that INH induces uncoupling of the mitochondrial membrane proton leak without promoting ATP synthesis and causes dysfunction in mitochondrial complexes I-IV, while ATP synthase activity remains unaffected. Additionally, we observed increased Ca^2+^ transfer from the ER to mitochondria (Fig. 4M). In support of mitochondrial morphology analysis, we detected an increase of mitoKeima-Red, a mitophagy marker, in the INH-treated groups (Fig. 4N).

These findings highlight the impact of INH on mitochondrial membrane integrity and functions, including impairment of mitochondrial complex I-IV, uncoupling of the mitochondrial membrane proton leak causing mitochondrial depolarization, and increased mitochondrial fission and mitophagy.

### ER stress is related to cellular ROS generation and altered redox balance during early INH exposure

To investigate if mitochondrial ROS is the critical factor impairing mitochondrial function and causing cellular oxidative stress, cellular ROS was monitored using cellROX and mitochondrial ROS using mitoSOX from 12 to 72 h after INH treatment. A significant increase in cellular ROS was observed at 12 h (Fig. 5A). In comparison, mitochondrial ROS levels increased only after 48 h (Fig. 5B). This suggests that INH initially increases cytosolic ROS, followed by a later increase in mitochondrial ROS, indicating that mitochondrial complex dysfunction may not be the primary cause of INH-induced cellular oxidative stress.

**Fig. 5.**
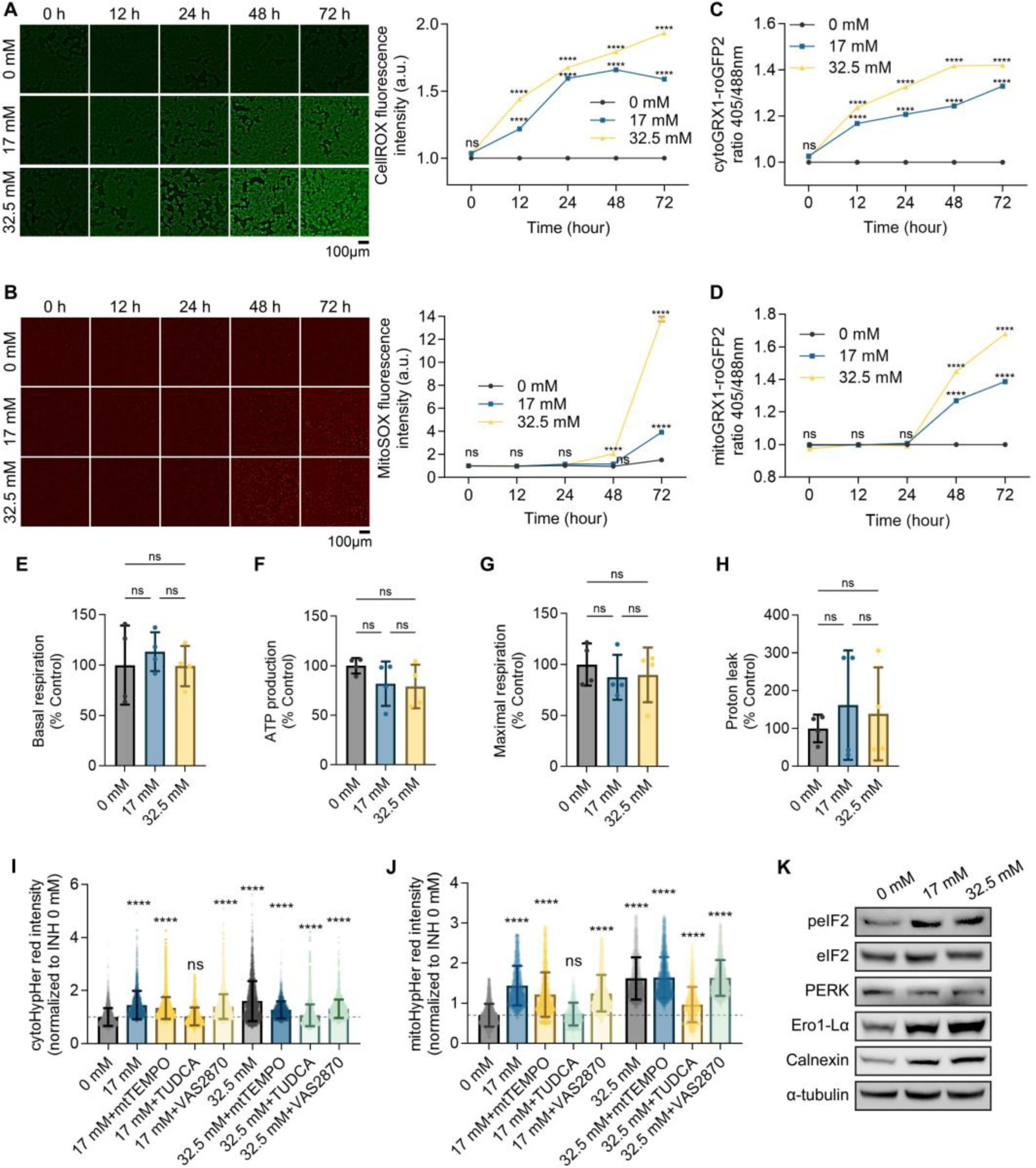
Cytosolic ROS levels rapidly increased after INH exposure, followed by a subsequent rise in mitochondrial ROS. (A-B) Time-dependent effects of INH treatment on cellular and mitochondrial ROS levels in HepG2 cells. (A) Representative images of HepG2 cells treated with 0 mM, 17 mM, and 32.5 mM INH for 0, 12, 24, 48, and 72 h, stained with CellROX Green to detect cellular ROS (left). Quantification of CellROX fluorescence intensity over time is shown on the right. Scale bar: 2 mm. (B) Representative images of HepG2 cells treated with 0 mM, 17 mM, and 32.5 mM INH for 0, 12, 24, 48, and 72 h, stained with MitoSOX Red to visualize mitochondrial ROS (left). The right panel shows the quantification of MitoSOX fluorescence intensity over time. Scale bar: 2 mm. (C) Quantification of the cytoGRX1-roGFP2 fluorescence intensity ratio (405/488 nm) in HepG2 cells treated with 0 mM, 17 mM, and 32.5 mM INH for 0, 12, 24, 48, and 72 h, indicating changes in cytosolic redox state (GSSG/GSH ratio) over time. (D) Measurement of the mitoGRX1-roGFP2 fluorescence intensity ratio (405/488 nm) in HepG2 cells treated with 0 mM, 17 mM, and 32.5 mM INH for 0, 12, 24, 48, and 72 h, showing alterations in the mitochondrial redox state (GSSG/GSH ratio) over time. (E-H) Mitochondrial oxygen consumption rate in HepG2 cells was determined using mitoXpress after 24-h INH treatment. The analysis includes (E) basal respiration rate, (F) ATP production rate, (G) maximal respiration rate, and (H) proton leak. (I) Cytosolic ROS levels at 24 h, determined using cytoHyPer-Red signals, in INH-treated cells alone or in combination with different compounds: mitoTEMPO 15 µM, TUDCA 1mM, and VAS2870 10 µM. (J) Mitochondrial ROS levels at 48 h, monitored using mitochondrial HyPer-Red, in cells treated with INH alone or in mitoTEMPO 15 µM, TUDCA 1mM, and VAS2870 10 µM. (K) Western blot analysis of the indicated proteins following INH treatment for 24 h. Data are displayed as means ± SD, based on analysis of more than 100 cells per group. Statistical differences between groups were determined using one-way ANOVA with Tukey’s post-test for multiple comparisons; n.s., not significant, ****P-value < 0.0001.

To further explore the impact of increased ROS on glutathione oxidation, the ratio of oxidized glutathione (GSSG) to reduced glutathione (GSH) was measured using cytoplasmic and mitochondrial ratio metric glutathione sensors (Fig. 5C-D) from 12 to 72 h after INH treatment. Supporting the ROS time-lapse data, cytosolic GSSG/GSH was significantly higher from 12 h, while it only markedly increased in mitochondria from 48 h. Additionally, measurements of OCR 24 h after INH exposure did not show significant changes in basal respiration rate, ATP production, maximal respiratory capacity, or proton leak in both the INH-treated and control groups (Fig. 5E-H). This suggests that mitochondrial dysfunction is not the primary cause of cellular oxidative stress triggered by INH.

To identify the primary source of cytosolic ROS early increase, inducing the antioxidant response at 24 h, we inhibited mitochondrial ROS, ER stress, and its associated ROS, and membrane-associated NADPH oxidases using mitoTEMPO, TUDCA, and VAS2870, respectively [24]. As illustrated in Figure 5I-J, the ER stress inhibitor TUDCA successfully reduced both cytoplasmic and mitochondrial ROS at 24 and 48 h after INH exposure, indicating that the ROS primarily originated from ER stress. We further observed a significant increase in the protein levels of peIF2, Ero1-Lα, and calnexin 24 h after INH treatment, as demonstrated by the immunoblotting analysis (Fig. 5K). This suggests that the PERK/eIF2 pathway is activated early during INH exposure, further connecting ER stress to the observed rise in ROS levels.

Collectively, these findings indicate that INH treatment primarily triggers cytosolic ROS induction related to ER stress. This induction is followed by a subsequent rise in mitochondrial ROS, which may result from the diffusion of ROS from the cytosol or mitochondrial complex dysfunction, ultimately altering glutathione oxidation.

### Nrf2 activation is the key antioxidant defense in response to INH-induced oxidative stress

In the presence of cellular ROS, transcription factor Nrf2 (also known as nuclear factor erythroid 2-related factor 2, NFE2L2) can be activated and translocated into the nucleus to enhance the expression of multiple antioxidant genes [25]. To assess the activation of Nrf2 triggered by INH, we examined the expression of multiple reported Nrf2 targets using quantitative PCR (qPCR), including *HMOX1*, *SLC7A11*, *GCLC*, and *NQO1* from 24 h to 72 h (Fig. 6 A-D). We observed the upregulation of mRNA expression of these Nrf2 targets in INH 32.5 mM treatment from 24 h and remarkably increased at 72 h in both 17 mM and 32.5 mM treatment groups. Furthermore, we observed that Nrf2 was increasingly localized to the nucleus 24 h after exposure to INH, indicating enhanced activation of Nrf2 signaling pathways at this time point (Fig. 6E).

**Fig. 6.**
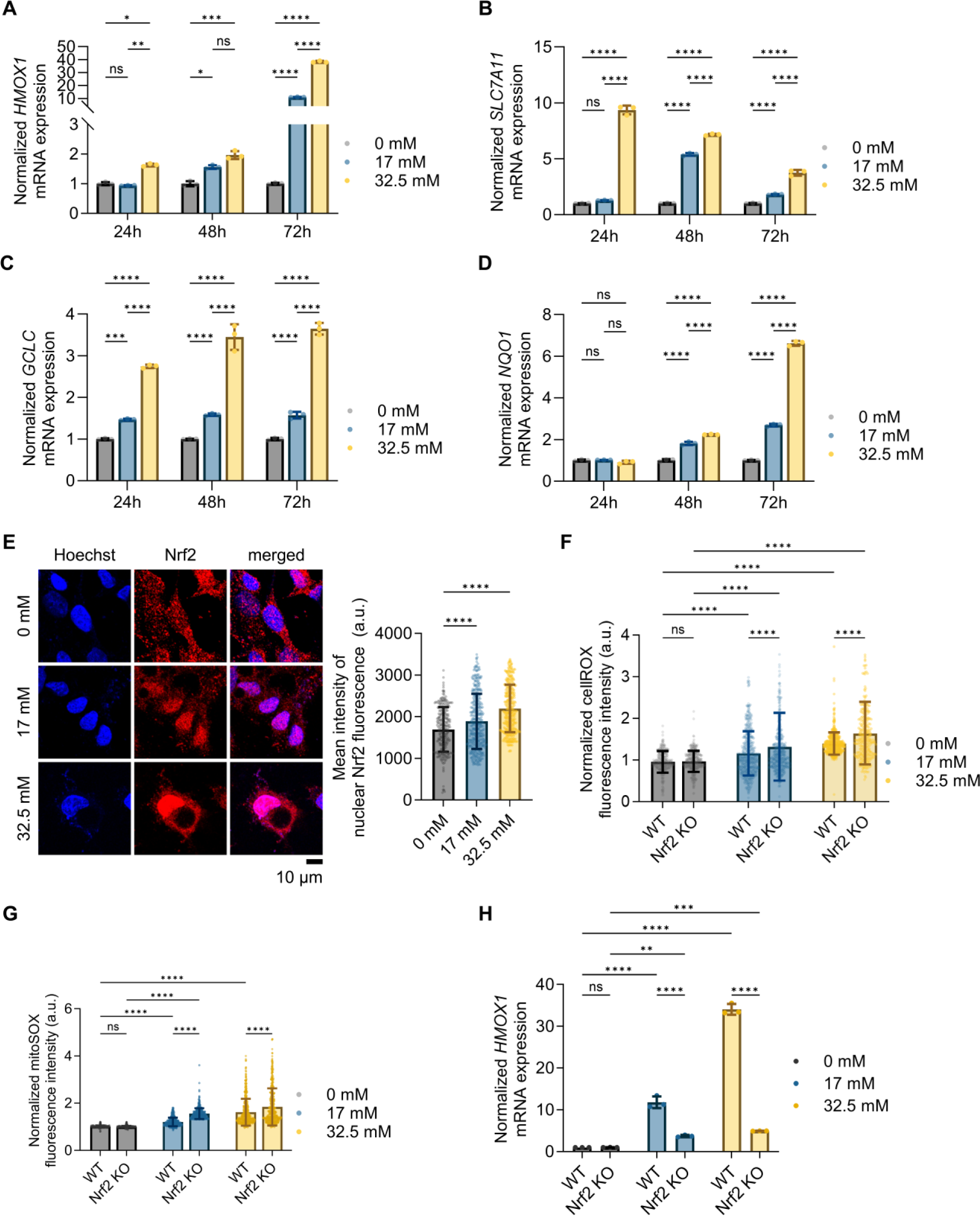
Oxidative stress triggered by INH exposure induces Nrf2 activation. (A-D) Normalized mRNA expression levels of Nrf2-targeted genes in HepG2 cells treated with 0 mM, 17 mM, and 32.5 mM INH for 24, 48, and 72 h. (A) *HMOX1*, (B) *SLC7A11*, (C) *GCLC,* and (D) *NQO1* mRNA expression measured using qPCR. (E) Representative immunofluorescence images of HepG2 cells treated with 0 mM, 17 mM, and 32.5 mM INH for 24 h, stained for Nrf2 (red) and Hoechst (blue). Merged images show nuclear localization of Nrf2 in INH-treated groups. Scale bar: 10 µm. The graph on the right quantifies the mean intensity of nuclear Nrf2 fluorescence. (F) Normalized CellROX fluorescence intensity in WT and Nrf2 KO HepG2 cells treated with 0 mM, 17 mM, and 32.5 mM INH for 72 h. (G) Quantification of mitochondrial ROS levels in WT and Nrf2 KO cells using MitoSOX fluorescence intensity after 72 h INH treatment. (H) *HMOX1* mRNA expression levels in WT and Nrf2 KO cells measured by qPCR. Data are shown as means ± SD. Statistical differences between groups were determined using one-way ANOVA with Tukey’s post-test for multiple comparisons; n.s., not significant, *P-value < 0.05, **P-value < 0.01, ***P-value < 0.001, ****P-value < 0.0001.

To further validate the role of Nrf2 in INH-induced oxidative stress, we generated Nrf2-KO cells using the CRISPR/Cas9 system, as shown in Supplementary Fig. S5. In these Nrf2-KO cells, we observed a significant increase in both cytosolic ROS (Fig. 6F) and mitochondrial ROS (Fig. 6G) when exposed to INH for 72 h compared to WT groups. This increase in ROS levels indicates heightened oxidative stress due to the absence of Nrf2. Moreover, we found that Nrf2 was essential for the upregulation of the antioxidant gene *HMOX1* in cells exposed to INH (Fig. 6H), further emphasizing the role of Nrf2 in regulating the cellular response to oxidative stress.

These findings demonstrate that Nrf2 signaling is crucial for cellular adaptation to INH-induced oxidative stress, highlighting its role in upregulating key antioxidant defenses to counteract ROS damage.

### Multi-omics unveils the critical role of Nrf2 in antioxidation defense and cell survival under INH exposure

We investigated the effects of Nrf2 KO on HepG2 cells treated with INH for 72 h. The Nrf2 KO cells demonstrated increased susceptibility to INH, exhibiting lower cell viability compared to WT HepG2 cells under the same INH concentration (Supplementary Fig. S6, Supplementary Fig. S7A). This data suggests that Nrf2 plays a protective role in mitigating INH-induced cytotoxicity.

We further investigated the molecular profile differences between Nrf2-KO and WT HepG2 cells under 72-hour INH treatment, finding substantial differences in their proteomics profiles, with 1745 differential proteins identified between the two cell types (Supplementary Fig. S7, Supplementary Fig. S7B, Supplementary Fig. S7C). Pathway enrichment analysis on these proteins revealed a disturbance in the apoptosis pathway, suggesting that KO cells were more vulnerable to INH treatments than WT cells (Supplementary Table S4). Furthermore, we observed perturbation in biological processes associated with mitochondrial function, including those related to the TCA cycle (“Citrate cycle (TCA cycle)”, “Pyruvate metabolism,” “Oxidative phosphorylation” and “Glycolysis / Gluconeogenesis”) and β-oxidation (“Fatty acid metabolism,” “Fatty acid degradation,” and “PPAR signaling pathway”). In addition, pathways related to ER stress (“Protein processing in endoplasmic reticulum” and “Protein export”) were shown to be changed (Supplementary Table S4). The GO:CC terms relevant to mitochondrial respiratory complexes and ER were found to be significant (Supplementary Table S5). These disturbances indicated the mitochondrial dysfunction and ER stress induced by INH treatment, which could be more severe in Nrf2-KO HepG2 cells. Moreover, compared to WT cells, KO-cells exhibited perturbations in multiple pathways related to antioxidative response. They included “Glutathione metabolism,” “Cysteine and methionine metabolism,” “Ferroptosis,” “HIF-1 signaling pathway,” “Selenocompound metabolism,” “Pentose phosphate pathway,” “Sulfur metabolism,” “Ascorbate and aldarate metabolism,” and “Glycine, serine and threonine metabolism” (Supplementary Table S4). Noticeably, except for the three former pathways, the six latter pathways were not significantly altered when comparing INH-treated WT cells with untreated WT cells. Given the loss of protection against oxidative stress from the Nrf2 signaling system, the newly founded significant pathways might represent alternative mechanisms that help KO cells manage oxidative stress. However, these mechanisms are insufficient to maintain cellular homeostasis. Besides, we also observed significant alterations in pathways related to endogenous cellular metabolism, i.e., amino acids metabolism, carbon metabolism, and lipid metabolism (Supplementary Table S4).

We captured a subtle difference in the metabolome between KO and WT cells under the same INH treatments with only 9 significantly differential metabolites (Supplementary Fig. S7D-F). The pathway analysis results showed metabolic perturbation related to redox balance, i.e., “Glutathione metabolism.” Indeed, we found that, under INH treatments, abundances of GSH and GSSG decreased in KO cells compared to WT cells (Supplementary Fig. S7F). We observed substantial alterations in the lipid profiles of KO compared to WT cells upon INH treatments, with 409 significantly differential lipids between these two types of cells (Supplementary Fig. S7G-H). It should be noted that basal levels of lipids in KO cells were substantially different from those of WT cells (Supplementary Fig. S7I). The levels of complex lipids such as glycerophospholipids, sphingolipids, and ceramide were significantly lower. The levels of lipid species involved in lipid droplet formation, regardless of INH treatment, were higher than WT. However, no significant differences were observed in the levels of droplet-related lipids between KO-cell groups. In agreement with our lipidomics data (Supplementary Fig. S7I), we observed similar patterns with insignificant changes in the number, size, and fluorescence intensity of lipid droplets between KO-cell groups (Supplementary Fig. S7J).

Collectively, our findings emphasized the central role of Nrf2 in protecting HepG2 cells against INH-induced stress. Additionally, we suggested alternative mechanisms that may be involved in cell defense against INH-induced oxidative stress besides Nrf2.

## Discussion

In this study, we extensively elucidated the mechanism by which INH induces hepatotoxicity in liver cells, expanding beyond the previously understood focus on mitochondrial oxidative stress and dysfunction (Fig. 7). Our findings reveal that INH-induced cellular ROS, associated with ER stress, occurs prior to the increase in mitochondrial ROS and subsequent mitochondrial dysfunction. Furthermore, prolonged INH exposure results in mitochondrial dysfunction related to the impairment of mitochondrial complexes I-IV and a proton leak in the mitochondrial membrane, without affecting mitochondrial ATP synthase. Additionally, INH prompted metabolic reprogramming in response to disruptions in the TCA cycle and beta-oxidation, leading to the accumulation of fatty acids and polyunsaturated fatty acids.

**Fig. 7.**
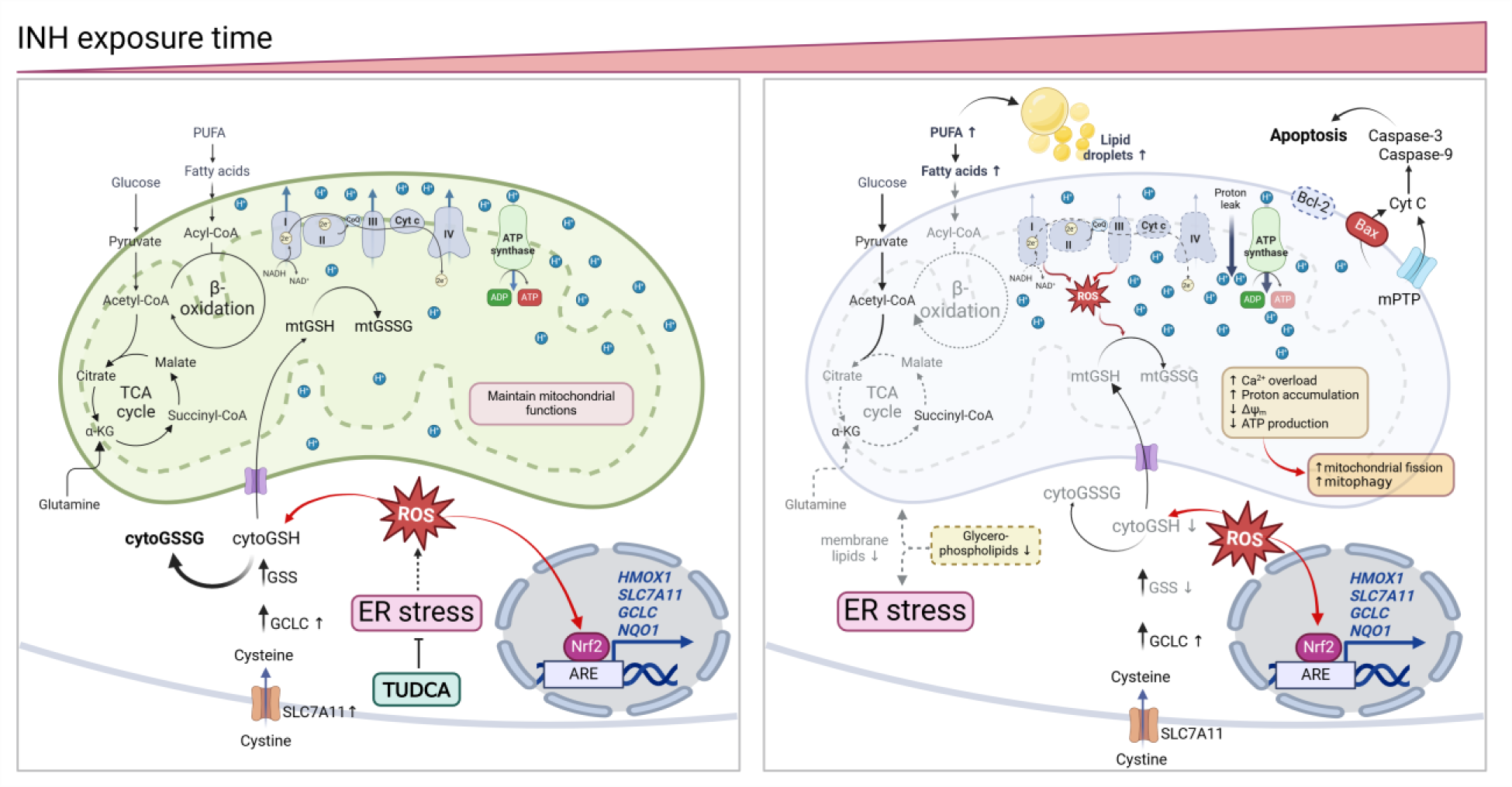
Mechanisms of INH-induced cellular oxidative stress and mitochondrial dysfunction in HepG2 cells. INH exposure initially induced cytosolic ROS rise, causing antioxidant responses including glutathione system (GSH/GSSG) and Nrf2-targeted antioxidant genes (*HMOX1*, *SLC7A11*, *GCLC*, *NQO1*), enhancing cellular defense mechanisms. Mitochondrial ROS has not risen during this period, and mitochondrial homeostasis is adequately maintained. By inhibiting ER stress during this stage using TUDCA, INH-induced cytosolic ROS can be eliminated. However, prolonged INH exposure results in antioxidant defense becoming overwhelmed, resulting in increased mitochondrial ROS in combination with mitochondrial complex I-IV defects and uncoupling mitochondrial membrane proton leaks that lead to mitochondrial Ca^2+^ overload, proton accumulation, mitochondrial depolarization, impaired ATP production, and insufficient TCA cycle. These changes resulted in mitochondrial fission, mitophagy and ultimately, apoptosis, as indicated by the activation of caspase-3 and caspase-9 and BAX translocation. In addition, there are elevated levels of complex lipids with polyunsaturated fatty acyl chains, fatty acids, and lipid droplet formation, indicating a global lipid alteration of the liver cell due to prolonged INH exposure.

We identified defects in mitochondrial complexes I-IV. On the other hand, an increased expression of ATP synthase (mitochondrial complex V) was found to be a compensatory mechanism by which the cell attempts to maintain ATP production. These results corroborate a previous study indicating that hydrazine, a major reactive metabolite of INH, can inhibit mitochondrial complex II activity [8]. In addition, INH induced a mitochondrial proton leak, supporting the “uncoupling to survive” hypothesis [26], which should be further verified. This mechanism generally exists to minimize oxidative damage by reducing the proton motive force and mitochondrial superoxide production [27]. Our proteomics data also revealed an increase in the protein abundance of aspartate-glutamate carriers (SLC25A12, SLC25A13), dicarboxylate carriers (SLC25A4, SLC25A5), and the phosphate carrier (SLC25A3) following INH treatment. Evidence supports that these members of mitochondrial solute carriers can facilitate mitochondrial proton leak [28].

Mitochondrial oxidative stress and dysfunction have long been regarded as the primary factors in INH-induced hepatotoxicity [7]. Our study provides evidence that mitochondrial superoxide levels increase concurrently with mitochondrial dysfunction after prolonged INH exposure. Interestingly, cellular ROS levels, but not the mitochondrial ROS levels, and the activation of the Nrf2 pathway were observed as early responses to INH treatment. In line with our findings, Chowdhury A *et al.* reported that INH 50 mg/kg alone induced hepatic oxidative stress in mice. However, they did not observe an increase in oxidative stress in the mitochondria of these mice [10]. Using TUDCA, an ER stress inhibitor, we found that ER stress and ER-associated ROS are the primary sources of cellular ROS elevation in early INH exposure time. However, TUDCA functions as a chemical chaperone, simultaneously mitigating ER protein misfolding and ER redox imbalance [29]. Inside the ER, ROS are mainly produced from the oxidative protein folding process, which is mediated by protein disulfide isomerase (PDIs) and ER oxidoreductin-1 (Ero1) proteins [30], and NOX4–a member of the NADPH oxidase (NOX) family [31]. Regarding the activation of PERK/eIF2 signaling, we also observed the upregulation of ER oxidoreductin-1 alpha (ERO1A) in early INH treatment. ERO1A generates hydrogen peroxide (H_2_O_2_) as a byproduct while reoxidizing PDI, transferring disulfide bonds (-S-S-) to newly synthesized proteins, which are essential for their correct folding. Moreover, hydrazine is a potent reducing agent that can donate its electrons to the disulfide bonds of proteins or GSSG, converting them into two free thiol groups (-SH) [32, 33]. Thus, INH or hydrazine may directly affect the ER protein unfolding/misfolding process and induce ER redox imbalance. Under ER stress, the accumulation of unfolded or misfolded proteins increases protein folding load, leading to excessive ROS (H_2_O_2_) production as a byproduct of disulfide bond formation. This process also depletes GSH, which is crucial to refolding misfolded proteins. Consequently, the overproduction of ROS and GSH depletion leads to oxidative stress [34, 35].

Upon INH exposure, the TCA cycle was impaired, partially reflected by the depletion of its intermediates. The mechanism of how INH impairs the TCA cycle is possibly due to the reaction between INH and keto acids, key intermediates of the TCA cycle [36]. In addition, a previous study showed that hydrazine can induce the depletion of TCA cycle intermediates [37]. Our data also suggested the disruption of β-oxidation, another critical process for maintaining cellular energy homeostasis. One potential mechanism for this disruption is the suppression of the carnitine shuttle process, which plays a vital role in transporting long-chain fatty acids to the mitochondrial matrix [38]. Specifically, the later step of transferring acylcarnitines to the mitochondrial matrix may be insufficient, as indicated by increments in the abundance of acylcarnitines with longer chains and fatty acids, as well as a decrease in the level of carnitine. The impairments of mitochondrial TCA cycle, β-oxidation, and oxidative phosphorylation may lead to energy homeostasis disorder.

Interestingly, the current study highlights several adaptations to the INH-induced energy crisis, implicating potential intervention strategies to prevent and ameliorate hepatotoxicity. We observed increased abundances of complex V protein subunits, TCA cycle enzymes, and β-oxidation enzymes, suggesting a compensatory effort to enhance ATP production as well as support energy and biosynthetic demands. Additionally, impairment of β-oxidation might lead to a deficiency in acetyl-CoA. Therefore, the increase in the conversion of pyruvate to acetyl-CoA may ensure consistent supply for the TCA cycle and lipid biosynthesis. Indeed, the expression levels of multiple enzymes involved in pyruvate metabolism were upregulated. Moreover, the abundances of CP1A (carnitine palmitoyltransferase 1A) and CPT2 (carnitine palmitoyltransferase 2), two main enzymes involved in carnitine-based fatty acids transport [39], were upregulated. This might be an effort to rescue the carnitine shuttle pathway to facilitate the β-oxidation process. Besides, lipid synthesis and turnover maintain membrane integrity and produce crucial signaling molecules. Together, these changes demonstrate a sophisticated cellular adaptation, enabling compensation for impaired mitochondrial function and ensuring cell survival and functionality.

Importantly, our study underscores the pivotal role of Nrf2 signaling in antioxidation defense and cellular survival in HepG2 cells subjected to INH treatment. Notably, cells lacking Nrf2 are particularly vulnerable to INH-induced oxidative stress, highlighting the critical importance of Nrf2 in cellular defense mechanisms. We identified alternative pathways involved in cellular antioxidant defense in Nrf2 KO conditions, such as selenocompound metabolism, the HIF-1 signaling pathway, and the pentose phosphate pathway. Selenocompound metabolism is critical for counteracting oxidative stress through selenoproteins like glutathione peroxidases (GPxs) and thioredoxin reductases (TrxRs), which use selenium to neutralize ROS, thus protecting cells from damage. Hypoxia-inducible factor-1 (HIF-1) regulates genes linked to oxidative stress, inflammation, and cell death [40], and is associated with liver toxicity in various contexts, including drug-induced liver injury [41]. A case-control study indicates that the HIF-1α gene is linked to a higher risk of anti-tuberculosis drug-induced liver injury (ATB-DILI) [42], and it has been shown that anti-TB drug-induced hepatocyte ferroptosis is controlled by the HIF-1α/SLC7A11/GPx4 signaling pathway [43]. Given the crucial roles of selenoproteins and HIF-1 signaling in antioxidant defenses, there is limited evidence on how these factors contribute to the progression of ATB-DILI, particularly INH-DILI.

Nrf2 plays a significant role in regulating lipid metabolism, especially in the context of GSH synthesis in the liver. According to a study by Asantewaa et al. (2024), GSH synthesis in the liver is essential for maintaining lipid abundance [44]. The repression of Nrf2 leads to increased lipid production and storage. In our study, Nrf2 KO cells displayed higher levels of lipid droplets and overall lipid storage compared to WT cells, regardless of INH treatments. Our data are consistent with the previous finding demonstrating that the repression of lipid storage and triglyceride levels is contingent on Nrf2 activation [44].

There are a few limitations and grounds for future studies. Even though widely used for toxicological investigation, molecular alterations of HepG2 cells may not fully represent the responses of normal liver cells. Additionally, our study helped uncover some potentially important questions regarding INH-induced toxicity for future investigations: (1) The detailed mechanism underlying INH-induced ER stress and the involvement of NOX4 in this molecular event, (2) the exact mechanisms by which INH or its metabolites impair mitochondrial complexes and the role of uncoupling proteins in mitochondrial dysfunction, and (3) the crosstalk between GSH and Nrf2 signaling in modulating cellular lipidome upon INH treatment. Future research should also validate our findings in animal or more physiologically relevant models, such as *in vitro* 3D hepatic microtissues, to better understand the *in vivo* relevance. Investigating the interactions between INH metabolites and mitochondrial complexes and exploring therapeutic strategies targeting Nrf2 and alternative antioxidant pathways could provide deeper insights and potential host-directed preventative therapy for INH-induced liver injury.

In conclusion, this study provides significant insights into the mechanisms of INH-induced hepatotoxicity, highlighting the complex interplay between ER stress, oxidative stress, mitochondrial dysfunction, and metabolic reprogramming. We demonstrated that ER stress is the primary source of ROS generation, initiating cellular oxidative stress in early INH exposure followed by mitochondrial oxidative stress in the later stages. The anti-ER-stress treatment can suppress the INH-induced ROS formation and protect cells against oxidative stress, which is the potential for developing preventative and alleviative therapy towards INH-induced liver injury. Our findings also emphasize the central role of Nrf2 signaling in the antioxidant response against INH-derived oxidative stress.

## Acknowledgment

Fig.7 was created using Biorender.com

## CRediT authorship contribution statement

**Truong Thi My Nhung**: Data curation, Methodology, Investigation, Formal analysis, Visualization, Validation, Writing – Original Draft, Writing – Review & Editing. **Nguyen Ky Phat**: Data curation, Methodology, Investigation, Formal analysis, Visualization, Validation, Writing – Original Draft, Writing – Review & Editing. **Trinh Tam Anh**: Methodology, Formal analysis, Visualization, Validation, Writing – Review & Editing. **Tran Diem Nghi**: Methodology, Formal analysis, Visualization, Validation, Writing – Review & Editing. **Nguyen Quang Thu**: Data curation, Formal analysis, Visualization, Validation, Writing – Review & Editing. **Ara Lee**: Methodology, Formal analysis, Software, Writing – Review & Editing. **Nguyen Tran Nam Tien**: Methodology, Formal analysis, Software, Writing – Review & Editing. **Nguyen Ky Anh**: Methodology, Formal analysis, Writing – Review & Editing. **Kimoon Kim:** Supervision, Methodology, Validation, Resources, Writing – Review & Editing. **Duc Ninh Nguyen:** Methodology, Validation, Writing – Review & Editing. **Dong Hyun Kim**: Supervision, Resources, Writing – Review & Editing, Funding acquisition. **Sang Ki Park**: Methodology, Investigation, Resources, Supervision, Writing – Review & Editing, Funding acquisition. **Nguyen Phuoc Long**: Conceptualization, Methodology, Investigation, Formal analysis, Validation, Resources, Supervision, Writing – Original draft, Writing – Review & Editing, Funding acquisition.

## Reference

1. World Health Organization Global tuberculosis report 2023. World Health Organization. 2023.

2. World Health Organization, WHO consolidated guidelines on tuberculosis: Module 4: Treatment - Drug-susceptible tuberculosis treatment. 2022: Geneva.

3. Forget E.J. and Menzies D. Adverse reactions to first-line antituberculosis drugs. Expert Opin Drug Saf. 2006;5(2):231–49.

4. Nahid P., Dorman S.E., Alipanah N., Barry P.M., Brozek J.L., Cattamanchi A., et al. Official American Thoracic Society/Centers for Disease Control and Prevention/Infectious Diseases Society of America Clinical Practice Guidelines: Treatment of Drug-Susceptible Tuberculosis. Clin Infect Dis. 2016;63(7):e147–e95.

5. Horsburgh C.R., Jr., Barry C.E., 3rd, and Lange C. Treatment of Tuberculosis. N Engl J Med. 2015;373(22):2149–60.

6. Nolan C.M., Goldberg S.V., and Buskin S.E. Hepatotoxicity associated with isoniazid preventive therapy: a 7-year survey from a public health tuberculosis clinic. JAMA. 1999;281(11):1014–8.

7. Boelsterli U.A. and Lee K.K. Mechanisms of isoniazid-induced idiosyncratic liver injury: emerging role of mitochondrial stress. J Gastroenterol Hepatol. 2014;29(4):678–87.

8. Lee K.K., Fujimoto K., Zhang C., Schwall C.T., Alder N.N., Pinkert C.A., et al. Isoniazid-induced cell death is precipitated by underlying mitochondrial complex I dysfunction in mouse hepatocytes. Free Radic Biol Med. 2013;65:584–94.

9. Ahadpour M., Eskandari M.R., Mashayekhi V., Haj Mohammad Ebrahim Tehrani K., Jafarian I., Naserzadeh P., et al. Mitochondrial oxidative stress and dysfunction induced by isoniazid: study on isolated rat liver and brain mitochondria. Drug Chem Toxicol. 2016;39(2):224–32.

10. Chowdhury A., Santra A., Bhattacharjee K., Ghatak S., Saha D.R., and Dhali G.K. Mitochondrial oxidative stress and permeability transition in isoniazid and rifampicin induced liver injury in mice. J Hepatol. 2006;45(1):117–26.

11. Zhang T., Ikejima T., Li L., Wu R., Yuan X., Zhao J., et al. Impairment of Mitochondrial Biogenesis and Dynamics Involved in Isoniazid-Induced Apoptosis of HepG2 Cells Was Alleviated by p38 MAPK Pathway. Front Pharmacol. 2017;8:753.

12. Schwab C.E. and Tuschl H. In vitro studies on the toxicity of isoniazid in different cell lines. Hum Exp Toxicol. 2003;22(11):607–15.

13. Shen C., Meng Q., Zhang G., and Hu W. Rifampicin exacerbates isoniazid-induced toxicity in human but not in rat hepatocytes in tissue-like cultures. Br J Pharmacol. 2008;153(4):784–91.

14. Cullinan S.B., Gordan J.D., Jin J., Harper J.W., and Diehl J.A. The Keap1-BTB protein is an adaptor that bridges Nrf2 to a Cul3-based E3 ligase: oxidative stress sensing by a Cul3-Keap1 ligase. Mol Cell Biol. 2004;24(19):8477–86.

15. Kobayashi A., Kang M.I., Okawa H., Ohtsuji M., Zenke Y., Chiba T., et al. Oxidative stress sensor Keap1 functions as an adaptor for Cul3-based E3 ligase to regulate proteasomal degradation of Nrf2. Mol Cell Biol. 2004;24(16):7130–9.

16. Zhang D.D. and Hannink M. Distinct cysteine residues in Keap1 are required for Keap1-dependent ubiquitination of Nrf2 and for stabilization of Nrf2 by chemopreventive agents and oxidative stress. Mol Cell Biol. 2003;23(22):8137–51.

17. Sasaki H., Sato H., Kuriyama-Matsumura K., Sato K., Maebara K., Wang H., et al. Electrophile response element-mediated induction of the cystine/glutamate exchange transporter gene expression. J Biol Chem. 2002;277(47):44765–71.

18. Jia Z.L., Cen J., Wang J.B., Zhang F., Xia Q., Wang X., et al. Mechanism of isoniazid-induced hepatotoxicity in zebrafish larvae: Activation of ROS-mediated ERS, apoptosis and the Nrf2 pathway. Chemosphere. 2019;227:541–50.

19. Ritchie M.E., Phipson B., Wu D., Hu Y., Law C.W., Shi W., et al. limma powers differential expression analyses for RNA-sequencing and microarray studies. Nucleic Acids Res. 2015;43(7):e47.

20. Liu P., Ewald J., Pang Z., Legrand E., Jeon Y.S., Sangiovanni J., et al. ExpressAnalyst: A unified platform for RNA-sequencing analysis in non-model species. Nat Commun. 2023;14(1):2995.

21. Pang Z., Lu Y., Zhou G., Hui F., Xu L., Viau C., et al. MetaboAnalyst 6.0: towards a unified platform for metabolomics data processing, analysis and interpretation. Nucleic Acids Res. 2024;52(W1):W398–W406.

22. Ramachandran A., Visschers R.G.J., Duan L., Akakpo J.Y., and Jaeschke H. Mitochondrial dysfunction as a mechanism of drug-induced hepatotoxicity: current understanding and future perspectives. J Clin Transl Res. 2018;4(1):75–100.

23. Bhadauria S., Mishra R., Kanchan R., Tripathi C., Srivastava A., Tiwari A., et al. Isoniazid-induced apoptosis in HepG2 cells: generation of oxidative stress and Bcl-2 down-regulation. Toxicol Mech Methods. 2010;20(5):242–51.

24. Romani P., Nirchio N., Arboit M., Barbieri V., Tosi A., Michielin F., et al. Mitochondrial fission links ECM mechanotransduction to metabolic redox homeostasis and metastatic chemotherapy resistance. Nat Cell Biol. 2022;24(2):168–80.

25. Tonelli C., Chio I.I.C., and Tuveson D.A. Transcriptional Regulation by Nrf2. Antioxid Redox Signal. 2018;29(17):1727–45.

26. Brand M.D. Uncoupling to survive? The role of mitochondrial inefficiency in ageing. Exp Gerontol. 2000;35(6-7):811–20.

27. Lambert A.J. and Brand M.D. Superoxide production by NADH:ubiquinone oxidoreductase (complex I) depends on the pH gradient across the mitochondrial inner membrane. Biochem J. 2004;382(Pt 2):511–7.

28. Skulachev V.P. Uncoupling: new approaches to an old problem of bioenergetics. Biochim Biophys Acta. 1998;1363(2):100–24.

29. Abdel-Ghaffar A., Elhossary G.G., Mahmoud A.M., Elshazly A.H.M., Hassanin O.A., Saleh A., et al. Potential prophylactic effect of chemical chaperones for alleviation of endoplasmic reticulum stress in experimental diabetic cataract. Bulletin of the National Research Centre. 2019;43(1):71.

30. Ramming T., Okumura M., Kanemura S., Baday S., Birk J., Moes S., et al. A PDI-catalyzed thiol-disulfide switch regulates the production of hydrogen peroxide by human Ero1. Free Radic Biol Med. 2015;83:361–72.

31. Laurindo F.R., Araujo T.L., and Abrahao T.B. Nox NADPH oxidases and the endoplasmic reticulum. Antioxid Redox Signal. 2014;20(17):2755–75.

32. Furst A., Berlo R.C., and Hooton S. Hydrazine as a Reducing Agent for Organic Compounds (Catalytic Hydrazine Reductions). Chemical Reviews. 1965;65(1):51–68.

33. Yew W.W., Chang K.C., and Chan D.P. Oxidative Stress and First-Line Antituberculosis Drug-Induced Hepatotoxicity. Antimicrob Agents Chemother. 2018;62(8).

34. Cao S.S. and Kaufman R.J. Endoplasmic reticulum stress and oxidative stress in cell fate decision and human disease. Antioxid Redox Signal. 2014;21(3):396–413.

35. Chen X., Shi C., He M., Xiong S., and Xia X. Endoplasmic reticulum stress: molecular mechanism and therapeutic targets. Signal Transduct Target Ther. 2023;8(1):352.

36. Li F., Miao Y., Zhang L., Neuenswander S.A., Douglas J.T., and Ma X. Metabolomic analysis reveals novel isoniazid metabolites and hydrazones in human urine. Drug Metab Pharmacokinet. 2011;26(6):569–76.

37. Bollard M.E., Keun H.C., Beckonert O., Ebbels T.M., Antti H., Nicholls A.W., et al. Comparative metabonomics of differential hydrazine toxicity in the rat and mouse. Toxicol Appl Pharmacol. 2005;204(2):135–51.

38. Longo N., Frigeni M., and Pasquali M. Carnitine transport and fatty acid oxidation. Biochim Biophys Acta. 2016;1863(10):2422–35.

39. Knottnerus S.J.G., Bleeker J.C., Wust R.C.I., Ferdinandusse S. L I.J., Wijburg F.A., et al. Disorders of mitochondrial long-chain fatty acid oxidation and the carnitine shuttle. Rev Endocr Metab Disord. 2018;19(1):93–106.

40. Kaminsky-Kolesnikov Y., Rauchbach E., Abu-Halaka D., Hahn M., Garcia-Ruiz C., Fernandez-Checa J.C., et al. Cholesterol Induces Nrf-2- and HIF-1alpha-Dependent Hepatocyte Proliferation and Liver Regeneration to Ameliorate Bile Acid Toxicity in Mouse Models of NASH and Fibrosis. Oxid Med Cell Longev. 2020;2020:5393761.

41. Chu Q., Gu X., Zheng Q., and Zhu H. Regulatory mechanism of HIF-1alpha and its role in liver diseases: a narrative review. Ann Transl Med. 2022;10(2):109.

42. Chong Y., Zhu H., Ren Q., Ma X., and Feng F. Interaction between the HIF-1alpha gene rs1957757 polymorphism and CpG island methylation in the promoter region is associated with the risk of anti-tuberculosis drug-induced liver injury in humans: A case-control study. J Clin Pharm Ther. 2022;47(7):948–55.

43. Liu Y., Chen W., Cen Y., Zhao X., Chen Z., Liang Y., et al. Hepatocyte ferroptosis contributes to anti-tuberculosis drug-induced liver injury: Involvement of the HIF-1alpha/SLC7A11/GPx4 axis. Chem Biol Interact. 2023;376:110439.

44. Asantewaa G., Tuttle E.T., Ward N.P., Kang Y.P., Kim Y., Kavanagh M.E., et al. Glutathione synthesis in the mouse liver supports lipid abundance through NRF2 repression. Nat Commun. 2024;15(1):6152.

