## Supplementary Material for "Endoplasmic reticulum stress inhibition preserves mitochondrial function and cell survival during early onset of isoniazid-induced oxidative stress" (from another version of this preprint: version 2, 2024-08-17)

### Supplementary Material and Methods

#### Antibodies

The study utilized various antibodies purchased from Cell Signalling Technology, including rabbit polyclonal anti-Bax (Cat#2772), rabbit monoclonal anti-Bcl-2 (D17C4)(Cat#3498), rabbit polyclonal anti-Caspase-9 (Cat#9502), rabbit monoclonal anti-cleaved Caspase-3 (Cat#9662), rabbit polyclonal anti-β-Actin (Cat#4967), rabbit monoclonal anti-Nrf2 (Cat#12721), rabbit monoclonal anti-VDAC (Cat#4661), rabbit polyclonal anti-eIF2α (Cat#9722), rabbit monoclonal anti-peIF2α (Cat#3597), rabbit monoclonal anti-PERK (Cat#5683), rabbit monoclonal anti-Calnexin (Cat#2679), rabbit polyclonal anti-Ero1-Lα (Cat#3264), rabbit polyclonal anti- α-tubulin (Cat#2144), HRP-conjugated horse anti-mouse IgG (Cat#7076), and HRP-conjugated goat anti-rabbit IgG (Cat#7074). Additional antibodies included mouse monoclonal anti-α-tubulin (Proteintech, Cat#66031-1-Ig) and mouse monoclonal anti-GAPDH (Santa Cruz, Cat#32233).

#### Chemicals

The study utilized various chemicals purchased from Sigma-Aldrich, including ATP (Cat#A2383), oligomycin (Cat#75351), FCCP (Cat#C2920), rotenone (Cat#R8875), and antimycin A from *Streptomyces sp.* (Cat#A8674), MitoTEMPO (Cat#SML0737), Tauroursodeoxycholic Acid Sodium Salt (TUDCA) (Cat#580549), VAS2870 (Cat#SML0273), Poly-D-lysine (Cat#P0899). Additional reagents included HBSS (Gibco, Cat#14175095), and Fluorescence Mounting Medium (Agilent Technologies, Cat#S302380-2).

#### Plasmids

TOM20-mScarlet was generated in-house. GCaMP6mt was obtained from S.K. Park [1]. The Mito-AT1.03 plasmid was a gift from Hiroyuki Noji to S.K. Park [2]. The pC1-HyPer-Red-C199S, (mito-SypHer Red), pC1-HyPer-Red and pC1-HyPer-Red-mito plasmids were gifts from Vsevolod Belousov (Addgene, #48252, #48249, #60247) [3]. The pLPXC cyto Grx1-roGFP2 (Addgene, #64975) and pLPXC mito Grx1-roGFP2 (Addgene plasmid #64977) plasmids were gifts from Tobias Dick [4]. pMitophagy Keima-Red mPark2 was purchased from MBL Life science, #AM-V0259M.

**ROS detection**

To assess cellular ROS and mitochondrial superoxide production, cells were treated with INH for various time points (0 to 72 h). Post-treatment, cells were incubated with either DHE (Cayman, 601290-96), 5 μM CellROX Green Reagent (ThermoFisher, C10444), or a combination of 1 μM MitoSOX Red (ThermoFisher, M36008) and 2.5 μM MitoTracker™ Deep Red FM (ThermoFisher, M22426) for 30 minutes, depending on the experiment's purpose. Imaging was performed using a confocal microscope microscope with a UPLSAPO 20×/0.75 NA objective. CellROX Green was excited at 488 nm and emitted at 525 nm. DHE and MitoSOX Red were excited at 510 nm and emitted at 580 nm. MitoTracker™ Deep Red FM was excited at 644 nm and emitted at 665 nm. MitoTracker signals were used to map and segment the mitochondria, providing a reference for accurately localizing and quantifying the MitoSOX Red signals. The fluorescence intensity was acquired and normalized to the control at each time point.

**Immunoblotting**

Cells were seeded in 6-well plates (6x10^5 cells/well). Following INH treatment, cell lysates were prepared with RIPA buffer and analyzed on 10% -12% SDS-PAGE gels. Proteins were transferred to PVDF membranes, blocked with 5% skim milk or BSA, and incubated with primary antibodies: Cleaved Caspase-3 (1:500), Caspase-9 (1:1000), Bax (1:1000), Bcl-2 (1:500), eIF2α (1:1000), peIF2α (1:500), PERK (1:1000), Calnexin (1:1000), Ero1-Lα (1:1000), and Tubulin (1:2000). After washing, secondary antibody (1:2000) was applied, and signals were developed with ECL solution and visualized using the ChemiDoc system.

**Laser scanning confocal microscopy**

HepG2 WT cells were cultured on poly-D-lysine-coated coverslips in 24-well plates at a density of $1.5 \times 10^5$ cells/ mL. For imaging purposes, cells were transfected with designated plasmids using Lipofectamine 2000 (Thermo Fisher Scientific) according to the manufacturer's instructions for 24 h, followed by incubation in culture media containing 17 mM and 32.5 mM INH for 72 h.

For mitochondrial morphology analysis, after INH treatment, cells expressing TOM20-mScarlet were fixed and imaged using an FV3000 confocal laser scanning microscope (Olympus) with a UPLSAPO 100X/1.4 NA objective. The excitation and emission wavelengths for mScarlet were 561 nm and 586 nm, respectively. The acquired images were deconvolved using the ClearView™ Accelerated Deconvolution algorithms in Imaris 9.8 software (Bitplane) for laser scanning confocal microscopy. Mitochondrial morphology was analyzed in at least 100 cells per condition, focusing on mitochondrial content, area, aspect ratio, and absolute number using a published ImageJ macro analysis method [5].

For tetramethyl rhodamine methyl ester (TMRM) imaging, cells after INH-exposure were incubated in 20 nM TMRM (Merck, T5428) in Hanks' Balanced Salt Solution (HBSS) supplemented with 25 mM HEPES, 2 mM $CaCl_2$, 1 mM $MgSO_4$, 4 mM $NaHCO_3$, and 30 mM D-glucose for 30 m at 37 °C. TMRM fluorescence intensity was recorded using a confocal microscope (UPLSAPO 20X objective / 0.75 NA) with 5 nM TMRM in the imaging solution. TMRM was excited at 543 nm, and emission was recorded at 573 nm. The total imaging time was 6 min with intervals of 15 s. Basal measurements were taken for 2 min, followed by 2-min recordings for each treatment (oligomycin, 2.5 μg/mL; FCCP, 5 μM). Data were collected using Cellsense software.

For mitochondrial pH measurement, cells were transfected with mitoSypHer Red (Addgene, 48252) following 72 h of INH treatment. Images were acquired using a confocal microscope equipped with a UPLSAPO 20X objective (0.75 NA), with excitation at 560 nm and emission at 645 nm. Image processing involved thresholding, single-cell segmentation, and intensity measurement using ImageJ.

For live mitochondrial ATP imaging, cells transfected with Mito-AT1.03 were exposed to INH for 72 h. Imaging was conducted using a confocal microscope with a UPLSAPO 20X objective (0.75 NA), using 405 nm for excitation. Emissions from CFP and YFP were simultaneously collected at 475 nm and 525 nm, respectively. The ratio of YFP to CFP fluorescence intensity was calculated to assess mitochondrial ATP levels in individual cells.

For calcium imaging, GCaMP6mt-expressing cells were treated with INH for 72 h. After treatment, the cells were imaged using a confocal microscope with a UPLSAPO 20×/0.75 NA objective. The medium was replaced with extracellular buffer (HBSS supplemented with 25 mM HEPES, 2 mM $CaCl_2$, 1 mM $MgSO_4$, 4 mM $NaHCO_3$, and 30 mM D-glucose. GCaMP6f was excited at 488 nm and emission was recorded at 509 nm. Imaging was conducted for 30 s to record basal intensity (Fo), followed by stimulation with 250 μM ATP to induce ER Ca2+ release. Total imaging time was 3 min with a 2-s interval. Intensity profiles of individual cells were obtained using Cellsense software (Olympus). After background subtraction, ΔF values were calculated as (F - Fo), where F represents the intensity of each time frame and Fo is the average intensity of 15 frames before stimulation, used for normalization.

For mitoKeimaRed imaging, cells were transfected with pMitophagy Keima-Red mPark2 (MBL Life science, AM-V0259M). After 24 hours of transfection, cells were treated with INH for 72 h. Imaging was conducted using a confocal microscope equipped with a UPLSAPO 20×/0.75 NA objective. MitoKeima-Red was excited at two wavelengths, 440 nm and 586 nm, and emission was detected at 620 nm. The fluorescence intensity ratios were calculated and normalized to the control at each time point to assess mitophagy.

To assess cellular ROS and mitochondrial superoxide production, cells were treated with INH for 0 to 72 h. Post-treatment, cells were incubated with 5 μM CellROX Green Reagent (ThermoFisher C10444) or 1 μM MitoSOX Red (ThermoFisher M36008) and 2.5 μM MitoTracker™ Deep Red FM (ThermoFisher M22426) for 30 m. Imaging was performed using a confocal microscope microscope with a UPLSAPO 20×/0.75 NA objective. CellROX Green was excited at 488 nm and emitted at 525 nm. MitoSOX Red was excited at 510 nm and emitted at 580 nm. MitoTracker™ Deep Red FM was excited at 644 nm and emitted at 665 nm. MitoTracker signals were used to map and segment the mitochondria, providing a reference for accurately localizing and quantifying the MitoSOX Red signals. The fluorescence intensity was acquired and normalized to the control at each time point.

For HyPer-Red imaging, cells were transfected with either pC1-HyPer-Red-mito (Addgene, #60247) or pC1-HyPer-Red (Addgene, #48249). Following 24 hours of transfection, cells were treated with INH for either 24 or 48 hours and then examined using a confocal microscope equipped with a UPLSAPO 20×/0.75 NA objective. HyPer-Red, whether cytosolic or mitochondrial, was excited at 510 nm and emitted at 580 nm. The fluorescence intensity was measured and normalized to the control at each respective time point.

To measure the cytosolic and mitochondrial GSSG/GSH ratio, cells were transfected with cyto Grx1-roGFP2 (Addgene, #64975) or mito Grx1-roGFP2 (Addgene, #64977) following INH treatment. Live cells were then imaged using a confocal microscope equipped with a UPLSAPO 20×/0.75 NA objective. Signals were obtained at 510 nm emission following excitation at 405 nm (oxidized) and 488 nm (reduced) at various time points from 0 to 72 h. The 405/488 ratio, representing the GSSG/GSH ratio, was calculated for each cell.

**Lipid droplet staining**

Cells were grown on poly-D-lysine-coated coverslips and treated with INH for 72 hours. After washing with PBS, cells were fixed with 4% paraformaldehyde for 10 minutes, washed again, and permeabilized with 0.1% Triton X-100 in PBS for 10 minutes. Cells were then incubated with 1 μM BODIPY (Thermo Fisher Scientific, #D3922) in PBS for 30 minutes at room temperature. Following three PBS washes, cells were mounted on slides using an antifade medium. Z-stack confocal images were captured with a FV3000 confocal laser scanning microscope (Olympus) using a UPLSAPO ×100/1.35 NA oil objective and a 488-nm laser. BODIPY signals were analyzed with Imaris Spots (Imaris, Bitplane) to quantify lipid droplets' area, number, and intensity, with fluorescence intensity normalized to the control.

**Immunocytochemistry**

For Nrf2 staining, cells were treated with INH for 24 hours, fixed with 4% paraformaldehyde for 10 minutes, and washed with PBS. Permeabilization was performed using 0.3% Triton X-100 in PBS for 15 minutes. Blocking was done with 5% goat serum in PBS containing 0.1% Triton X-100 for 1 hour, followed by overnight incubation at 4°C with Nrf2 antibodies (CST, #12721). After washing, Alexa Fluor 568-conjugated secondary antibody (Molecular Probes, #A-11011) was applied for 1 hour at room temperature. Cells were then stained with Hoechst 33342 (Thermo Fisher, #H3570) for 10 minutes and mounted with antifade medium. Confocal images were captured using a FLUOVIEW FV3000 microscope (Olympus) with a UPLSAPO ×60/1.35 NA oil objective. Hoechst was detected at

405 nm and Nrf2 at 568 nm. The Hoechst signal segmented nuclei, enabling measurement of Nrf2 intensity within the nuclear region, which was normalized to the control.

**Cellular metabolite extraction and metabolome data acquisition**

WT or Nrf2 KO HepG2 cells were seeded in 6-well plates at a density of $6 \times 10^5$ cells per well. After 72 h of treatment with INH, the cultured media was removed, followed by washing twice using pre-warmed PBS (Gibco, 10010023). Subsequently, the 6-well plate was plated on dry ice for further steps of metabolite extraction. In brief, the cells were scraped off the dish using a cell scraper (Fischer, 1159T48) and quenched using 1000 μL of six internal standards (ISs) "premixed" methanol (MeOH) (Sigma, 1060352500) at -80$^o$C. The solution was then vortexed briefly following a homogenization step using the Bead Ruptor Elite (Omni, 19-042E) (two times for 15 s at 5 m/s) with cooling on dry ice for 2 min between each homogenization. After a centrifugation step (for 10 min at 4 $^o$C and 16,000 *rcf*), the supernatant was collected and dried by $N_2$ flow at room temperature. The dried extracts were stored at -80$^o$C until further analysis. For subsequent analysis, the dried extracts were reconstituted with 100 μL of 50% MeOH in water (*v/v*) and vortexed for 30 s. The mixtures were then centrifuged for 5 min at 4°C and 16,000 *rcf*. From each sample, 70 μL of the supernatant was transferred to vials for LC-MS data acquisition, while 20 μL was reserved for post-extraction pooled quality control (QC) preparation. The samples were kept at 4 $^o$C in an autosampler during analysis.

Chromatographic separation was performed using the Shimadzu Nexera LC system (Kyoto, Japan). The characteristics of detected ions were captured with the X500R Quadrupole Time-of-Flight mass spectrometer (Q-ToF MS), which is equipped with a Turbo V™ ion source and Twin Spray probe (SCIEX, MA, USA). Data acquisition utilized an information-dependent acquisition mode. Injection volumes of 5 μL for both positive ion mode (ESI+) and negative ion mode (ESI-) were employed. To ensure consistent data quality, mass calibration was conducted after every six injections using the X500R's built-in calibrant delivery system (CDS). The fast LC gradient conditions using the Acquity UPLC HSS T3 column (50 x 2.1 mm, 1.8 µm) coupled to an Acquity UPLC HSS T3 VanGuard pre-column (5 mm × 2.1 mm; 1.8 µm) (Waters, Milford, MA, USA) were applied follow the methods by Cajka *et al*. [6], and MS settings followed protocols described in our previous works [7-9].

**Cellular lipid extraction and lipidome data acquisition**

The lipid extraction process followed our previously established protocol with slight modifications [10]. WT or Nrf2 KO HepG2 cells were seeded in 6-well plates at a density of 6 × $10^5$ cells per well. After 72 h of treatment with INH, the cultured media was removed following by 2 times washing using pre-warmed PBS (Gibco, 10010023). Subsequently, the 6-well plate was plated on dry ice for further steps of lipidomic extraction. The mixture of two IS solutions (1:1, *v/v*), including LIPIDOMIX® Mass Spec Standard and Deuterated Ceramide Mass Spec Standard (Avanti Polar Lipids, Alabama, USA) was dissolved in MeOH. The cells were scraped off the dish using a cell scraper and quenched using 250 µL of IS "premixed" MeOH (Sigma, 1060352500) at -80$^o$C. The solution was then homogenized using the Bead Ruptor Elite (Omni, 19-042E) (two times for 15 s at 5 m/s) with cooling on dry ice for 2 min between each homogenization. The homogenized solution was then transferred to a new 2 mL Eppendorf tube. Subsequently, 850 µL of pre-cooled (-20°C) methyl *tert*-butyl ether was added, vortexed for 60 s, and incubated on a shaker at 1,500 *rpm* and 4°C for 1 h. After adding 210 µL of pre-cooled (4°C) water, the mixture was vortexed again for 60 s and incubated on the shaker for an additional 15 min at the same conditions. The mixture was centrifuged for 10 min at 4 °C and 16,000 *rcf* . Next, 400 µL × 2 of the upper layer were transferred to the Eppendorf tube and evaporated using $N_2$ flow at room temperature. The resulting dry extracts were stored at -80°C until further analysis. Dried lipid extracts were resuspended in 100 μL of MeOH:toluene (9:1, *v/v*) followed by brief vortex, and centrifugation for 5 min at 4 °C and 16,000 *rcf* . 70 μL of the supernatant was transferred to the vial for subsequent instrumental analysis, and 20 μL of each sample was taken to make the post-extraction pooled QC samples. The samples were stored at 4°C in an autosampler during analysis.

For data acquisition, the Agilent UPLC Infinity II 1290 system (Agilent, USA) coupled with the X500R Q-ToF MS with a Turbo V™ ion source and Twin Spray probe (SCIEX, MA, USA) were utilized. For lipids separation, we used an Acquity BEH C18 column (2.1 × 50 mm, 1.7 µm) and an Acquity VanGuard BEH C18 pre-column (2.1 × 5 mm, 1.7 µm) (Waters, Miford, MA, USA) with gradient elution as previously detailed [11]. The injection volumes were optimized for each ion mode: 2 μL for ESI+, and 3 μL for ESI-, respectively. The data acquisition and tandem MS parameters were followed our previous studies. Mass calibration was performed via the CDS system after every sixth injection.

**Cellular protein extraction and proteome profiling data acquisition**

For proteomics analysis, HepG2 cells were seeded at a density of $1.5 \times 10^7$ cells per T75 flask for each experimental group. After a 24-h recovery period, cells were treated with various concentrations of INH (IC0, IC20, IC50), with the INH-supplemented culture medium being refreshed every 24 h. Following 72 h of treatment, cells were lysed using 2% SDS in LC-grade water containing protease inhibitor cocktails (1×). Cell lysis was facilitated using a tip sonicator (3 sets, 50% amplitude, 1 second with 15 pulses per set), and the lysates were centrifuged at 12,000 *rcf* and 25°C to remove debris. The resulting lysate was incubated with cold acetone (-20°C) for protein precipitation at -20°C for 2 h. The mixture was centrifuged for 15 min at 4°C and 13,000 *rcf*, the supernatant was discarded, and the precipitate was incubated in cold acetone (-20°C) with 10% TBS buffer for 1 h at -20°C. After another centrifugation for 15 min at 4°C and 13,000 *rcf*, the precipitate was air-dried for 10 min. The protein pellet was denatured overnight in 8 M urea in 50 mM ammonium bicarbonate (ABC) buffer, followed by sonication (3 sets, 50% amplitude, 1 s with 10 pulses per set). Protein concentration was determined using the bicinchoninic acid (BCA) assay (Thermo Fisher Scientific).

For reduction, 20 µg of proteins in 8 M urea in 50 mM ABC buffer were treated with DTT to a final concentration of 10 mM for 1 h at 37°C. Alkylation was performed by adding IAA to a final concentration of 40 mM for 1 h at 37°C. The solution was diluted to 1 M urea with ABC buffer (50 mM), supplemented with $CaCl_2$ to a final concentration of 1 mM, and digested with trypsin-TPCK (20:1 protein-to-trypsin ratio) at 37°C for 4 h. The digested solution was centrifuged for 15 min at 13,000 *rcf*, desalted using Pierce™ C18 Spin Tips (Thermo Scientific #87784), dried under vacuum overnight, and re-dissolved in LC-grade water with 0.1% TFA for LC-MS/MS analysis.

A Q-Exactive Plus mass spectrometer (Thermo Fisher Scientific) with an Ultimate 3000 nanoLC pump and autosampler was used to conduct NanoLC-MS/MS. An Easy Spray column (Thermo Fisher Scientific, #ES803A, C18, 2 µm, 100 Å, 500 mm length, 75 µm diameter) was used. The injection volume was set at 6 µL with a flow rate of 300 nL/min. The mobile phases were (A) 0.1% formic acid in water and (B) 0.1% formic acid in acetonitrile, with a gradient program set from 10% to 90% B over a total of 130 min. A spray voltage of 1.5 kV was applied, scanning m/z ranges from 350–1800, selecting the top 10 precursor ions for MS/MS scans with a normalized collision energy of 27 eV.

**Omics data processing, data treatment, and identification**

Regarding metabolomics and lipidomics, we utilized MS-DIAL version 4.9.0 to process the raw data (.wiff files), optimizing the parameters for each column and ion mode.

For MS/MS spectra inspection in metabolomics, we used the IRCCS Istituto Giannina Gaslini-Mass Spectra Library [12] and other MS-DIAL libraries. Additionally, we established in-house libraries using the Mass Spectrometry Metabolite Library of Standards (msmls-lot-230-01, Sigma Aldrich) and the Bile Acid/Carnitine/Sterol Metabolite Library of Standards (bacsmls-lot-205-36, Sigma Aldrich) to enhance the confidence of metabolite annotation. For lipidomics data processing, we utilized the MS-DIAL inbuilt library and Fiehn's lab lipidomics library [13, 14] for manually inspecting MS/MS spectra in lipid identification [7, 11]. After data processing, the aligned data were exported and treated using MetaboAnalyst 6.0. Features with ≥ 50% missing data were removed, and imputation was performed using k-nearest neighbors (k-NN) algorithms. Features with a relative standard deviation of more than 25% in the pooled QC groups were also filtered out. Consequently, the data in 2 ion modes of metabolomics and lipidomics were mixed. Only highly confident annotated metabolites and lipids were subjected to subsequent downstream analyses.

Regarding the proteomic data, Proteome Discoverer 2.3 (PD 2.3, Thermo Fisher Scientific) for peak list creation from recorded fragmentation spectra was used to process raw MS data files. Automated database searching was performed using SEQUEST against the UniprotKB/SwissProt *Homo sapiens* proteome database, with a precursor mass tolerance of 10 ppm, a fragment ion mass tolerance of 0.02 Da, and trypsin specificity allowing up to two missed cleavages. Peptides were identified if marked as 'Master' in PD 2.3. A reverse decoy database search established a false discovery rate (FDR) of less than 1% at both peptide and protein levels. Label-free quantification, averaging the relative peak intensities of identified peptides, was applied to quantify protein. The label-free peptide-level proteomics data was processed following the protocol of Hutchings *et al* [15]. In brief, peptide features that did not match a master protein, features corresponding to contaminant proteins, features with missing quantitative information, features linked to multiple protein groups, features with ranks that were not 1, and features that were not annotated as “unambiguous” were filtered out. Peptide features with more than 20% missing values were also removed. Missing values were imputed using the k-NN algorithm.

**Omics data normalization, exploratory analysis, and visualization**

Proteomics, metabolomics, and lipdidomics data were median normalized and log-transformed. The log-transformed normalized data were scaled before performing the principal component analysis (PCA) and heatmap visualization. Data exploration and visualization was conducted using MetaboAnalyst 6.0 [16] and R version 4.3.2. Heatmaps were visualized using *Complexheatmap* package version 2.20.0 [17] unless stated otherwise, and volcano plots were visualized using *EnhancedVolcano* package version 1.22.0.

**Functional analysis**

Differential proteins were submitted to over-representation analysis (ORA) performed using *clusterProfiler* package version 4.12.0 [18]. The Kyoto Encyclopedia of Genes and Genomes (KEGG) [19] and Gene Ontology Cellular Component (GO:CC) knowledgebases were used. A false discovery rate (FDR) via the Benjamini-Hochberg procedure of 0.05 was used as the significance threshold. KEGG pathway analysis using the list of significantly differential metabolites was performed using MetaboAnalyst 6.0 [16]. The significant pathways were determined based on the P-value threshold 0.05 and the significant hits > 1. For lipidomics, significantly differential lipids were subjected to the lipid ontology (LION) enrichment analysis via the “LION-PCA heatmap” module [20].

**Supplementary Figures**

**Supplementary Fig. S1. Differential protein analysis results of WT HepG2 exposing to INH.** (A) WT – 17 mM INH vs. WT – 0 mM INH. (B)  WT – 32.5 mM INH vs. WT – 0 mM INH. Abbreviation: WT, wild-type; INH, isoniazid; FDR, false discovery rate.

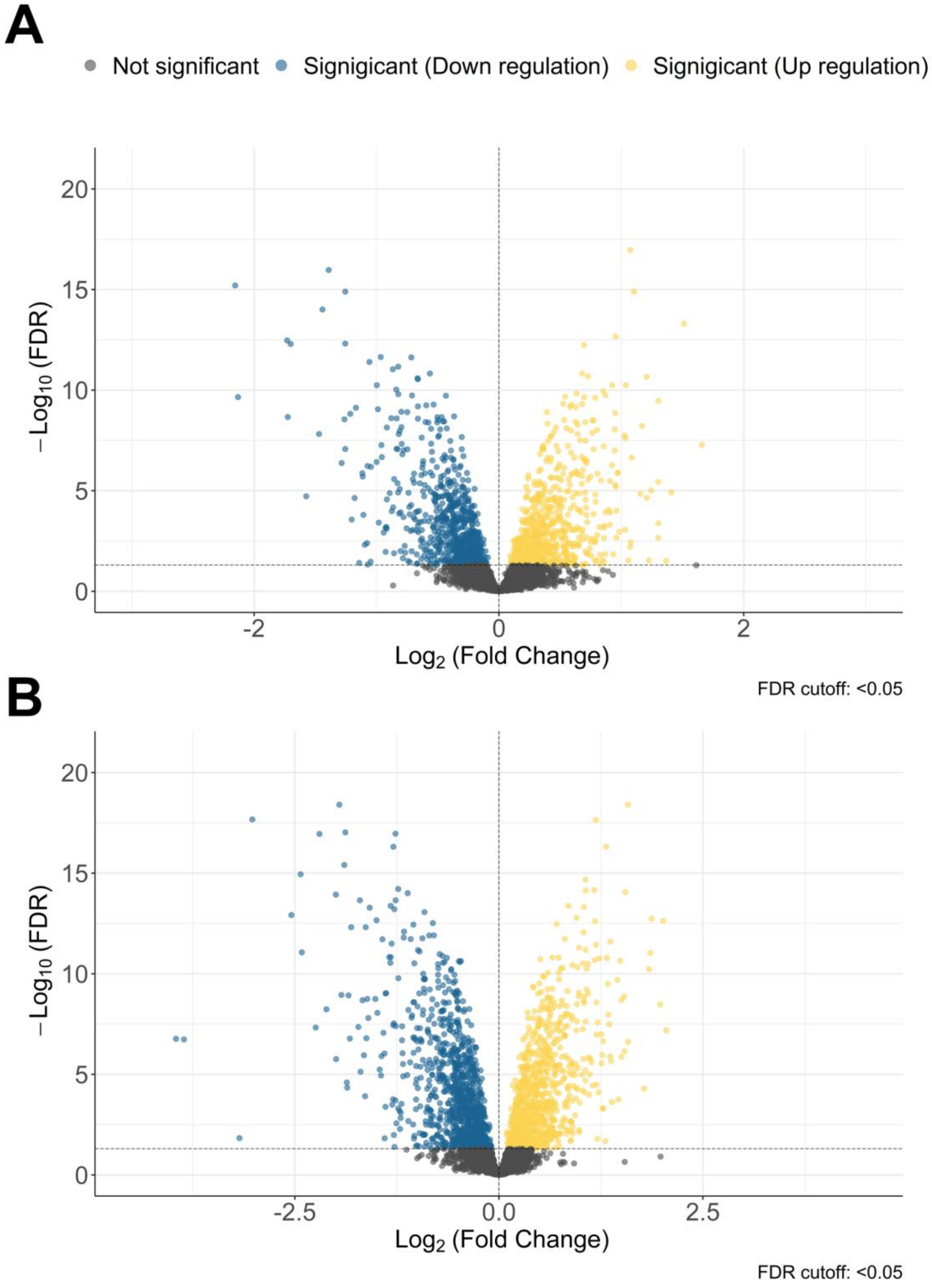

**Supplementary Fig. S2. Heatmap of significantly differential proteins between INH-treated and untreated WT HepG2 cells enriched into significant GO:CC terms.** (A) GO:CC terms related to endoplasmic reticulum. (B) GO:CC terms related to mitochondrial respiratory complex. Abbreviation: GO:CC, gene ontology (cellular component); WT, wild-type; FDR, false discovery rate.

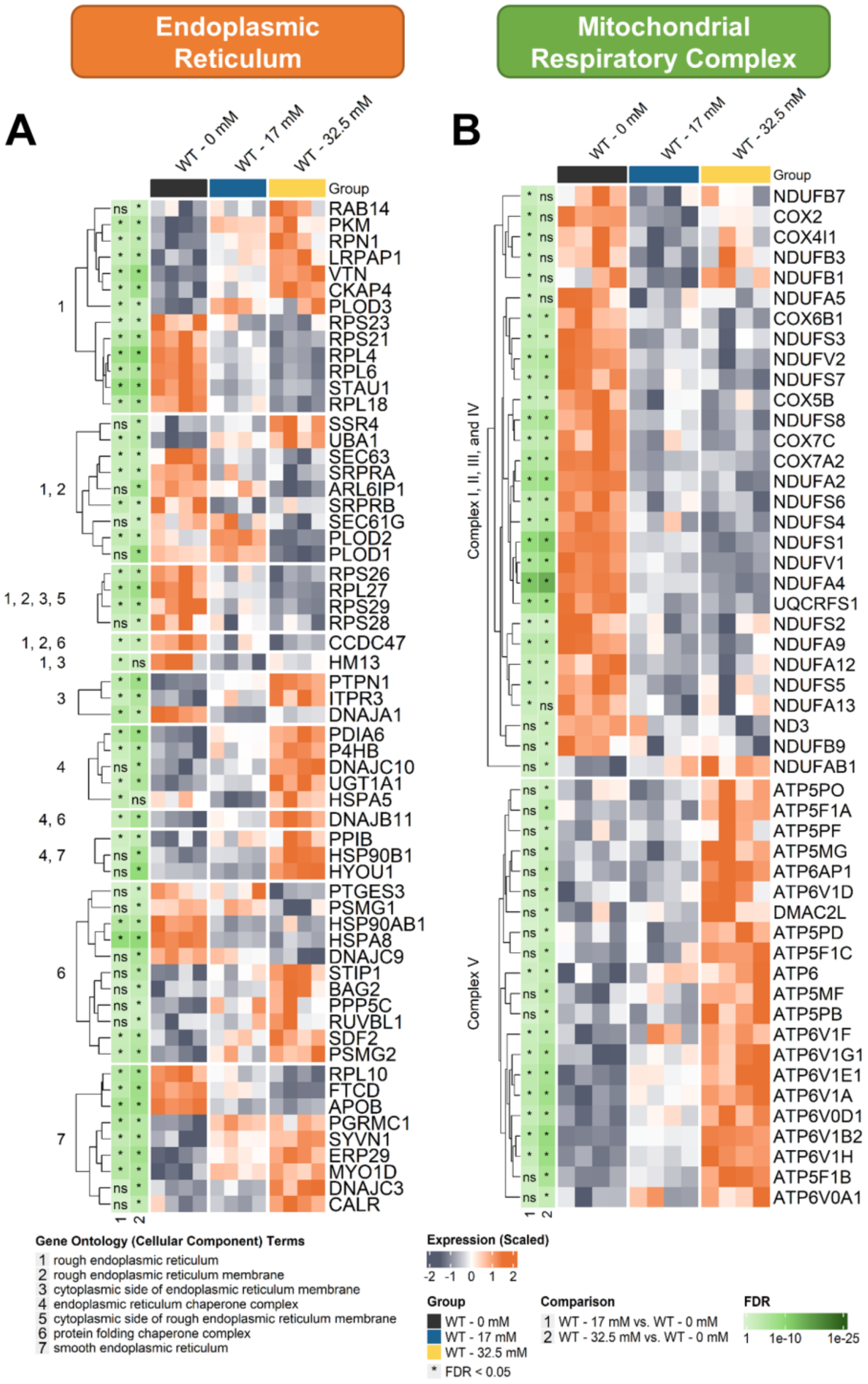

**Supplementary Fig. S3. Differential metabolite analysis and differential lipids analysis results of WT HepG2 cells exposing to INH.** (A) Metabolomics, WT – 17 mM INH vs. WT – 0 mM INH. (B) Metabolomics, WT – 32.5 mM INH vs. WT – 0 mM INH. (C) Lipidomics, WT – 17 mM INH vs. WT – 0 mM INH. (D) Lipidomics, WT – 32.5 mM INH vs. WT – 0 mM INH. Abbreviation: WT, wild-type; FDR, false discovery rate.

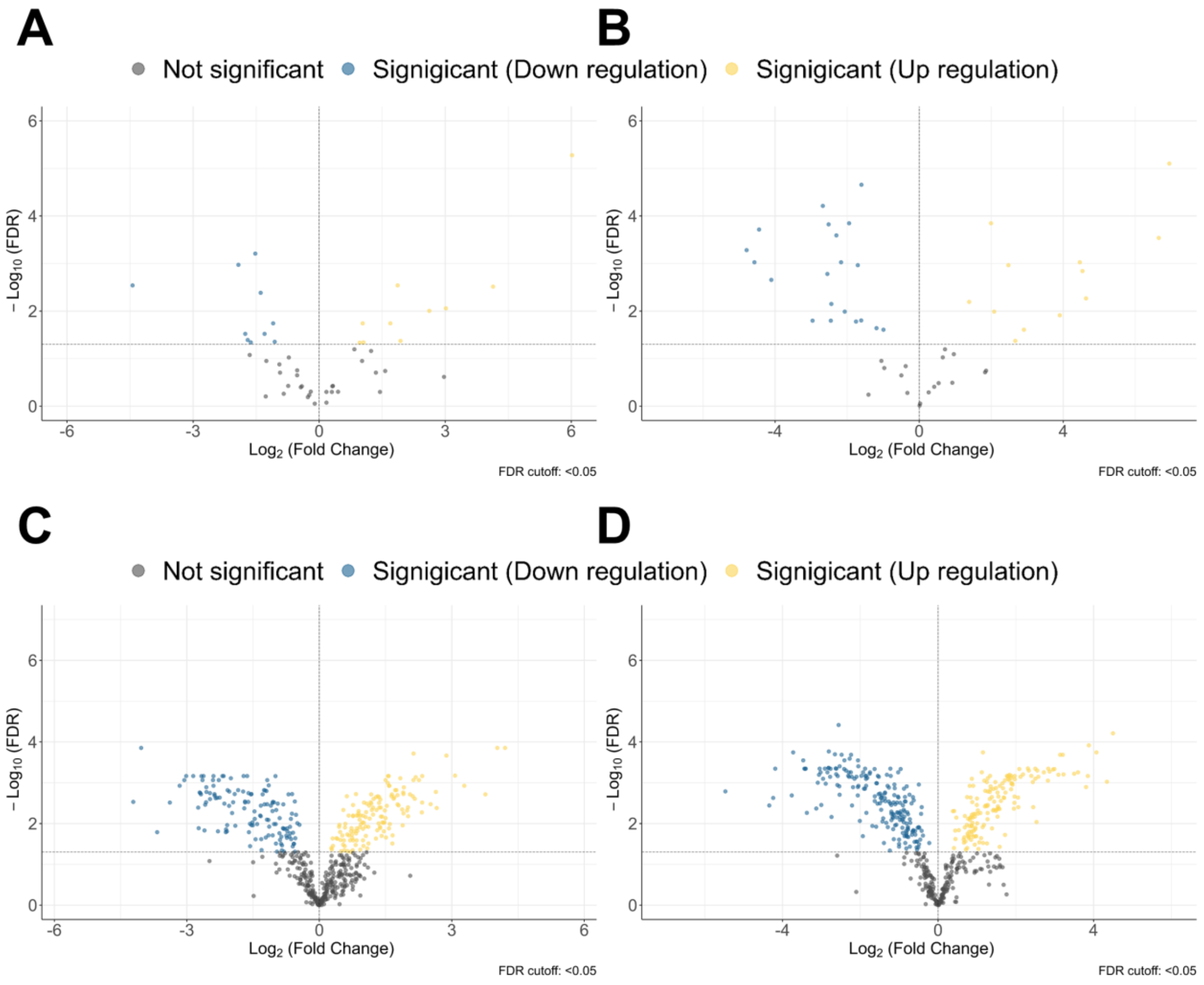

**Supplementary Fig. S4. Heatmaps of significantly differential lipids between INH-treated and untreated WT HepG2 cells.** Abbreviation: WT, wild-type; FDR, false discovery rate; FA, Fatty acid; NAE, N-acyl ethanolamines; DG, Diacylglycerol; DG(O-), Ether-linked diacylglycerol; TG, Triacylglycerol; TG(O-), Ether-linked triacylglycerol; PC, Phosphatidylcholine; PC(O-), Ether-linked phosphatidylcholine; PE, Phosphatidylethanolamine; PE(O-), Ether-linked phosphatidylethanolamine; BMP, Bismonoacylglycerophosphate; CL, Cardiolipin; LPC, Lysophophatidylcholine; LPC(O-), Ether-linked lysophosphatidylcholine; LPE, Lysophosphatidylethanolamine; LPE(O-), Ether-linked lysophosphatidylethanolamine; PG, Phosphatidylglycerol; PI, Phosphatidylinositol; PS, Phosphatidylserine; Cer, Ceramide; SM, Sphingomyelin; CE, Cholesteryl ester; CoQ, Coenzyme Q.

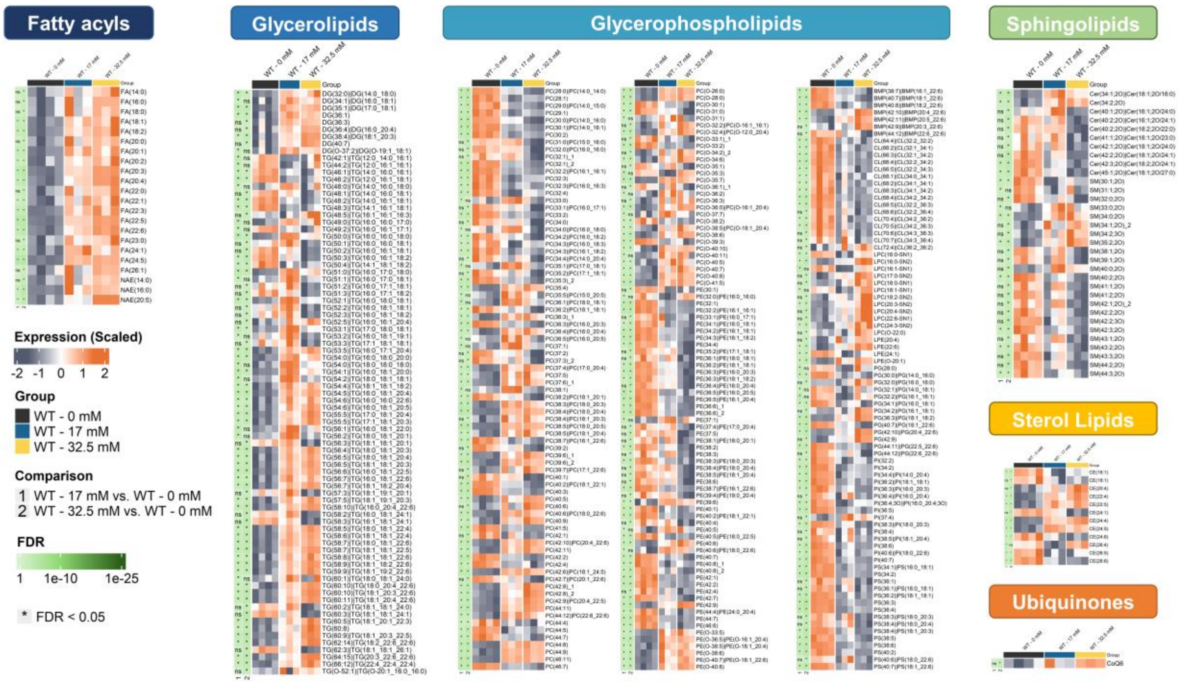

**Supplementary S5. Generation of Nrf2-knockout HepG2 cells using CRISPR-Cas9 system.** (A). Illustration of Nrf2-knockout process using CRISPR-Cas9 system. (B). Knockout of Nrf2 in HepG2 cell mediated by CRISPR Cas9.

A

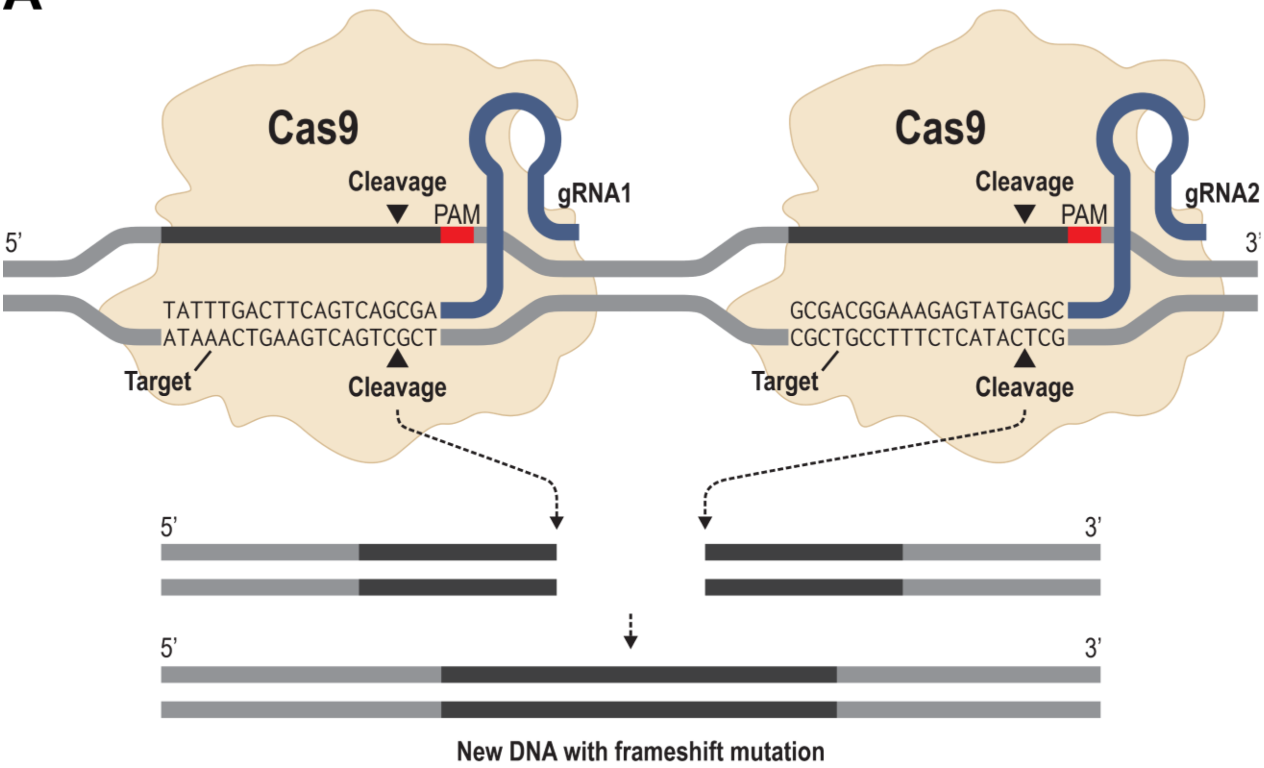


B

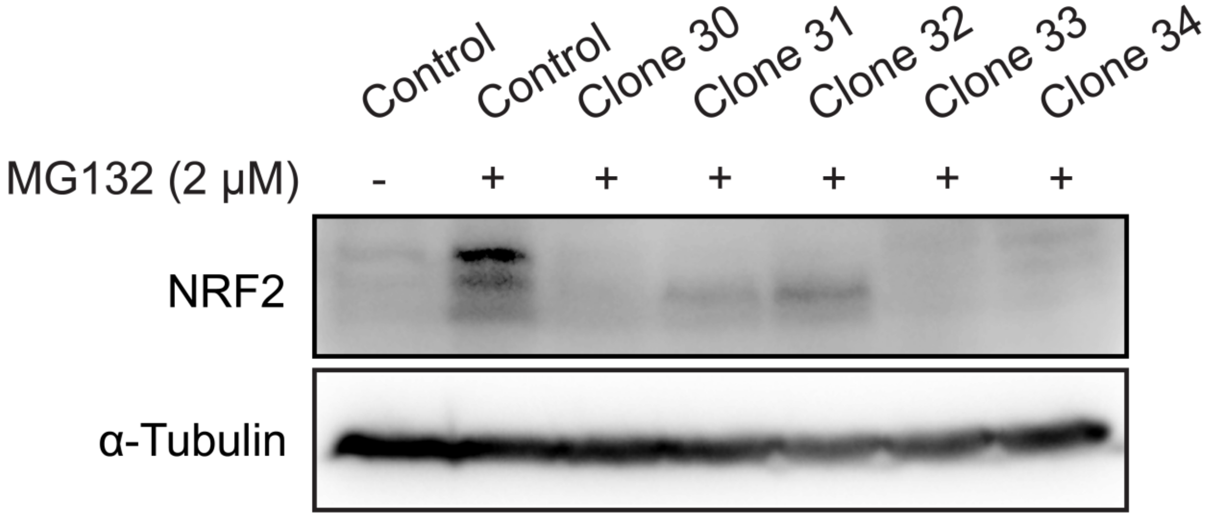

**Supplementary Fig S6. Dose response curve of Nrf2-KO HepG2 treated with different concentration of INH (0, 8.5, 17, 32.5, 50 mM).**

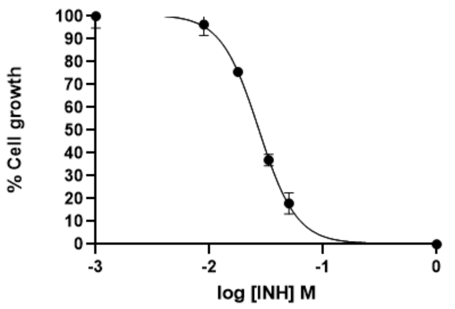

**Supplementary Fig S7. Multi-omics analysis of differences between Nrf2-KO and WT HepG2 cells upon INH treatments.** (A) Cell viability in Nrf2-KO and WT HepG2 cells treated with different INH concentrations. (B) Principal component analysis score plots of proteomics profiles of Nrf2-KO and WT HepG2 cells upon INH treatments. (C) Differential protein analysis results of Nrf2-KO versus WT HepG2 cells upon INH treatments. (D) Principal component analysis score plots of metabolomics profiles of Nrf2-KO and WT HepG2 cells upon INH treatments. (E) Differential metabolite analysis results of Nrf2-KO versus WT HepG2 cells upon INH treatments. (F) Heatmap of significantly differential metabolites between Nrf2-KO and WT HepG2 cells upon INH treatments. (G) Principal component analysis score plots of lipidomics profiles of Nrf2-KO and WT HepG2 cells upon INH treatments. (H) Differential lipids analysis results of Nrf2-KO versus WT HepG2 cells upon INH treatments. (I) Principal component analysis heatmap of lipid ontology enrichment analysis of significantly differential lipids between Nrf2-KO and WT HepG2 cells upon INH treatments. (J) Representative images of lipid droplets in Nrf2 KO HepG2 cells treated with 0 mM, 17 mM, and 32.5 mM INH, stained with BODIPY 493/503. Scale bar: 10 μm. Quantification includes the relative number, mean area, mean intensity of lipid droplets per cell. Data are presented as means ± SD. Statistical significance was determined using one-way ANOVA with Tukey's post-test for multiple comparisons; n.s., not significant, **P-value < 0.01. Abbreviation: WT, wild-type; KO, knockout.

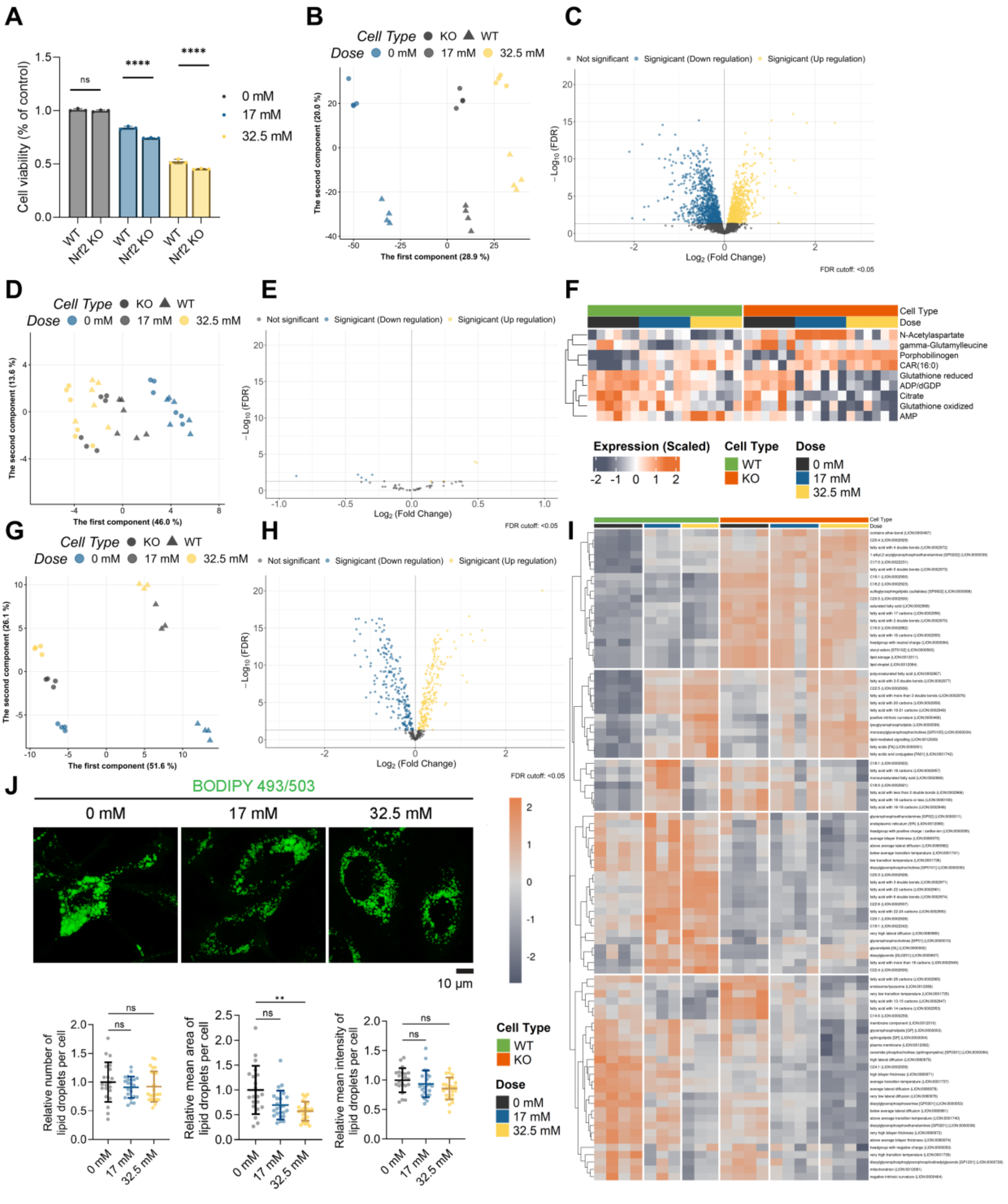
A
Cell viability (% of control)
1.5
1.0
0.5
0.0
ns
****
****
WT
Nrf2 KO
0 mM
17 mM
32.5 mM
B
Cell Type KO WT
Dose 0 mM 17 mM 32.5 mM
The second component (20.0 %)
The first component (28.9 %)
C
Not significant
Signigicant (Down regulation)
Signigicant (Up regulation)
−Log10 (FDR)
Log2 (Fold Change)
FDR cutoff: <0.05
D
Cell Type KO WT
Dose 0 mM 17 mM 32.5 mM
The second component (13.6 %)
The first component (46.0 %)
E
Not significant
Signigicant (Down regulation)
Signigicant (Up regulation)
−Log10 (FDR)
Log2 (Fold Change)
FDR cutoff: <0.05
F
Cell Type
Dose
N-Acetylaspartate
gamma-Glutamylleucine
Porphobilinogen
CAR(16:0)
Glutathione reduced
ADP/dGDP
Citrate
Glutathione oxidized
AMP
Expression (Scaled)
Cell Type
WT
KO
Dose
0 mM
17 mM
32.5 mM
G
Cell Type KO WT
Dose 0 mM 17 mM 32.5 mM
The second component (26.1 %)
The first component (51.6 %)
H
Not significant
Signigicant (Down regulation)
Signigicant (Up regulation)
−Log10 (FDR)
Log2 (Fold Change)
FDR cutoff: <0.05
I
Cell Type
Dose
J
BODIPY 493/503
0 mM
17 mM
32.5 mM
10 μm
Relative number of lipid droplets per cell
Relative mean area of lipid droplets per cell
Relative mean intensity of lipid droplets per cell
ns
**
Cell Type
WT
KO
Dose
0 mM
17 mM
32.5 mM

**Supplementary Tables**

**Supplementary Table S1. Significant pathways resulted from over-representation analysis of significantly differential proteins between isoniazid-treated and untreated wild-type HepG2 cells.**

| **KEGG Pathway** | **WT – 17 mM vs. WT – 0 mM** | | **WT – 32.5 mM vs. WT – 0 mM** | |
|---|---|---|---|---|
| | **FDR** | **Number of Enriched Proteins** | **FDR** | **Number of Enriched Proteins** |
| Ribosome | 3.59E-36 | 72 | 1.38E-35 | 80 |
| Coronavirus disease - COVID-19 | 5.29E-18 | 78 | 1.24E-13 | 83 |
| Spliceosome | 2.16E-12 | 46 | 6.82E-16 | 59 |
| Carbon metabolism | 2.94E-11 | 45 | 1.46E-11 | 53 |
| Lysosome | 2.63E-07 | 38 | 2.24E-04 | 37 |
| Biosynthesis of amino acids | 4.36E-07 | 28 | 6.11E-07 | 32 |
| Protein processing in endoplasmic reticulum | 1.73E-06 | 44 | 7.42E-08 | 56 |
| Oxidative phosphorylation | 8.83E-06 | 37 | 2.13E-07 | 48 |
| Chemical carcinogenesis - reactive oxygen species | 8.83E-06 | 54 | 7.49E-05 | 62 |
| Fatty acid degradation | 2.21E-05 | 19 | 9.80E-07 | 24 |
| Glycolysis / Gluconeogenesis | 3.18E-05 | 24 | 5.79E-04 | 25 |

| | | | | |
|---|---|---|---|---|
| Citrate cycle (TCA cycle) | 3.18E-05 | 15 | 1.30E-06 | 19 |
| Amino sugar and nucleotide sugar metabolism | 1.23E-04 | 18 | 2.72E-04 | 20 |
| Valine, leucine and isoleucine degradation | 2.02E-04 | 18 | 1.67E-08 | 28 |
| Pyruvate metabolism | 2.02E-04 | 18 | 1.39E-03 | 19 |
| Biosynthesis of cofactors | 2.54E-04 | 35 | 1.06E-05 | 46 |
| Aminoacyl-tRNA biosynthesis | 4.27E-04 | 15 | 1.28E-02 | 14 |
| Arginine and proline metabolism | 5.46E-04 | 17 | 7.42E-03 | 17 |
| Ribosome biogenesis in eukaryotes | 6.90E-04 | 19 | 3.14E-04 | 23 |
| Pentose and glucuronate interconversions | 7.19E-04 | 13 | - | - |
| Biosynthesis of nucleotide sugars | 7.74E-04 | 14 | 9.27E-04 | 16 |
| Diabetic cardiomyopathy | 7.90E-04 | 45 | 4.23E-03 | 52 |
| Fatty acid metabolism | 1.39E-03 | 18 | 6.11E-07 | 28 |
| Thermogenesis | 2.54E-03 | 45 | 4.67E-03 | 54 |
| Glutathione metabolism | 3.52E-03 | 17 | 2.05E-04 | 23 |
| Ascorbate and aldarate metabolism | 3.56E-03 | 11 | - | - |
| Parkinson disease | 3.56E-03 | 51 | 2.33E-03 | 64 |

| | | | | |
|---|---|---|---|---|
| Ferroptosis | 4.10E-03 | 14 | 1.40E-02 | 15 |
| Folate biosynthesis | 8.31E-03 | 9 | - | - |
| Nucleocytoplasmic transport | 1.11E-02 | 21 | 2.14E-07 | 37 |
| Non-alcoholic fatty liver disease | 1.13E-02 | 31 | - | - |
| PPAR signaling pathway | 1.65E-02 | 19 | - | - |
| Alanine, aspartate and glutamate metabolism | 1.77E-02 | 12 | 2.47E-03 | 16 |
| Other glycan degradation | 1.80E-02 | 7 | - | - |
| Galactose metabolism | 2.28E-02 | 10 | 4.07E-02 | 11 |
| Glyoxylate and dicarboxylate metabolism | 2.28E-02 | 10 | 5.37E-03 | 13 |
| Peroxisome | 2.31E-02 | 18 | 1.82E-03 | 25 |
| Prion disease | 2.31E-02 | 49 | 1.09E-02 | 63 |
| Huntington disease | 2.66E-02 | 50 | 9.85E-03 | 65 |
| Pentose phosphate pathway | 3.69E-02 | 9 | - | - |
| N-Glycan biosynthesis | 3.77E-02 | 12 | - | - |
| Histidine metabolism | 3.78E-02 | 7 | - | - |
| One carbon pool by folate | 4.54E-02 | 7 | 4.88E-02 | 8 |

| | | | | |
|---|---|---|---|---|
| 2-Oxocarboxylic acid metabolism | 4.63E-02 | 9 | 7.49E-05 | 16 |
| Various types of N-glycan biosynthesis | 4.63E-02 | 10 | - | - |
| Propanoate metabolism | - | - | 2.73E-06 | 19 |
| DNA replication | - | - | 9.86E-06 | 18 |
| Cysteine and methionine metabolism | - | - | 6.32E-05 | 23 |
| Mismatch repair | - | - | 2.24E-04 | 12 |
| Nucleotide metabolism | - | - | 5.59E-04 | 27 |
| mRNA surveillance pathway | - | - | 5.59E-04 | 27 |
| Butanoate metabolism | - | - | 1.82E-03 | 13 |
| Nucleotide excision repair | - | - | 3.01E-03 | 19 |
| Cholesterol metabolism | - | - | 5.51E-03 | 18 |
| Base excision repair | - | - | 6.80E-03 | 15 |
| Purine metabolism | - | - | 7.05E-03 | 32 |
| Fatty acid elongation | - | - | 1.33E-02 | 10 |
| Vibrio cholerae infection | - | - | 1.77E-02 | 16 |
| RNA degradation | - | - | 1.82E-02 | 20 |

| | | | | |
|---|---|---|---|---|
| Lipoic acid metabolism | - | - | 1.94E-02 | 8 |
| Tryptophan metabolism | - | - | 2.21E-02 | 14 |
| Starch and sucrose metabolism | - | - | 4.07E-02 | 11 |

**WT:** Wild-type

**FDR**: False discovery rate

**KEGG:** Kyoto Encyclopedia of Genes and Genomes

**Supplementary Table S2. Significant gene ontology (cellular component) terms resulted from over-representation analysis of significantly differential proteins between isoniazid-treated and untreated wild-type HepG2 cells.**

| Terms | WT – 17 mM vs. WT – 0 mM | | WT – 32.5 mM vs. WT – 0 mM | |
|---|---|---|---|---|
| | FDR | Number of Enriched Proteins | FDR | Number of Enriched Proteins |
| cytosolic large ribosomal subunit | 4.02E-24 | 40 | 4.41E-23 | 44 |
| small ribosomal subunit | 1.48E-21 | 42 | 1.18E-17 | 43 |
| cytosolic small ribosomal subunit | 2.19E-20 | 33 | 4.49E-19 | 36 |
| small-subunit processome | 5.19E-16 | 34 | 4.72E-16 | 39 |
| rough endoplasmic reticulum | 4.73E-08 | 22 | 4.46E-09 | 27 |
| eukaryotic translation initiation factor 3 complex | 1.32E-04 | 10 | 2.42E-03 | 9 |
| preribosome, large subunit precursor | 1.82E-04 | 9 | 6.20E-07 | 13 |
| mitochondrial respiratory chain complex I | 1.82E-04 | 19 | 1.53E-02 | 17 |
| NADH dehydrogenase complex | 1.82E-04 | 19 | 1.53E-02 | 17 |
| respiratory chain complex I | 1.82E-04 | 19 | 1.53E-02 | 17 |
| spliceosomal snRNP complex | 2.58E-04 | 17 | 1.72E-06 | 24 |
| eukaryotic 48S preinitiation complex | 4.31E-04 | 9 | 5.05E-04 | 10 |

| | | | | |
|---|---|---|---|---|
| preribosome, small subunit precursor | 5.04E-04 | 7 | 1.78E-03 | 7 |
| Sm-like protein family complex | 6.14E-04 | 19 | 5.25E-06 | 27 |
| eukaryotic 43S preinitiation complex | 6.60E-04 | 9 | 8.85E-04 | 10 |
| exon-exon junction complex | 7.14E-04 | 9 | 4.01E-03 | 9 |
| translation preinitiation complex | 7.14E-04 | 9 | 1.06E-03 | 10 |
| small nuclear ribonucleoprotein complex | 8.58E-04 | 17 | 1.18E-05 | 24 |
| precatalytic spliceosome | 1.28E-03 | 13 | 8.04E-07 | 21 |
| rough endoplasmic reticulum membrane | 1.94E-03 | 9 | 5.72E-06 | 14 |
| nucleoid | 1.98E-03 | 14 | 1.04E-05 | 21 |
| mitochondrial nucleoid | 1.98E-03 | 14 | 1.04E-05 | 21 |
| vacuolar proton-transporting V-type ATPase, V1 domain | 2.63E-03 | 6 | 1.66E-03 | 7 |
| proton-transporting V-type ATPase, V1 domain | 2.63E-03 | 6 | 1.66E-03 | 7 |
| U2-type precatalytic spliceosome | 2.67E-03 | 12 | 1.69E-06 | 20 |
| U2 snRNP | 3.93E-03 | 8 | 2.99E-04 | 11 |
| eukaryotic translation initiation factor 4F complex | 7.26E-03 | 7 | 2.08E-02 | 7 |
| cytoplasmic side of endoplasmic reticulum membrane | 7.26E-03 | 7 | 2.08E-02 | 7 |

| | | | | |
|---|---|---|---|---|
| paraspeckles | 7.69E-03 | 5 | 1.60E-02 | 5 |
| CRD-mediated mRNA stability complex | 9.64E-03 | 5 | 4.34E-03 | 6 |
| Cajal body | 9.64E-03 | 14 | 2.78E-02 | 15 |
| Prp19 complex | 1.03E-02 | 6 | 6.75E-03 | 7 |
| 90S preribosome | 1.04E-02 | 8 | 1.23E-03 | 11 |
| U2-type prespliceosome | 1.04E-02 | 7 | 1.03E-02 | 8 |
| prespliceosome | 1.04E-02 | 7 | 1.03E-02 | 8 |
| endoplasmic reticulum chaperone complex | 1.04E-02 | 6 | 1.86E-03 | 8 |
| mitochondrial respiratory chain complex IV | 1.05E-02 | 8 | - | - |
| flotillin complex | 1.19E-02 | 6 | - | - |
| respiratory chain complex IV | 1.53E-02 | 8 | - | - |
| RNA cap binding complex | 1.53E-02 | 7 | 4.31E-02 | 7 |
| replisome | 1.54E-02 | 7 | 1.57E-02 | 8 |
| sno(s)RNA-containing ribonucleoprotein complex | 1.54E-02 | 6 | 7.97E-04 | 9 |
| vacuolar proton-transporting V-type ATPase complex | 2.04E-02 | 7 | 6.73E-03 | 9 |
| proton-transporting two-sector ATPase complex, catalytic domain | 2.04E-02 | 6 | 3.46E-05 | 11 |

| | | | | |
|---|---|---|---|---|
| t-UTP complex | 2.27E-02 | 3 | - | - |
| cytoplasmic side of rough endoplasmic reticulum membrane | 2.27E-02 | 3 | 4.56E-03 | 4 |
| peroxisomal membrane | 2.43E-02 | 14 | 2.08E-02 | 17 |
| microbody membrane | 2.43E-02 | 14 | 2.08E-02 | 17 |
| B-WICH complex | 2.60E-02 | 4 | 1.04E-02 | 5 |
| nuclear replication fork | 2.67E-02 | 9 | - | - |
| proton-transporting V-type ATPase complex | 2.67E-02 | 7 | 3.57E-03 | 10 |
| podosome | 2.67E-02 | 10 | - | - |
| organellar small ribosomal subunit | 2.67E-02 | 7 | - | - |
| mitochondrial small ribosomal subunit | 2.67E-02 | 7 | - | - |
| eukaryotic translation initiation factor 3 complex, eIF3m | 2.98E-02 | 4 | - | - |
| vesicle tethering complex | 3.10E-02 | 13 | - | - |
| aminoacyl-tRNA synthetase multienzyme complex | 3.25E-02 | 5 | - | - |
| nuclear replisome | 3.48E-02 | 6 | 2.96E-02 | 7 |
| mCRD-mediated mRNA stability complex | 3.77E-02 | 4 | - | - |
| tertiary granule lumen | 3.90E-02 | 12 | 1.28E-02 | 16 |

| | | | | |
|---|---|---|---|---|
| peroxisomal matrix | 4.05E-02 | 12 | - | - |
| microbody lumen | 4.05E-02 | 12 | - | - |
| cytoplasmic ubiquitin ligase complex | 4.05E-02 | 5 | - | - |
| Derlin-1 retrotranslocation complex | 4.60E-02 | 4 | - | - |
| Hrd1p ubiquitin ligase ERAD-L complex | 4.98E-02 | 3 | - | - |
| U12-type spliceosomal complex | 4.98E-02 | 6 | 5.12E-03 | 9 |
| proton-transporting two-sector ATPase complex | - | - | 1.94E-06 | 21 |
| U2-type catalytic step 1 spliceosome | - | - | 1.43E-04 | 8 |
| catalytic step 1 spliceosome | - | - | 1.43E-04 | 8 |
| U2-type catalytic step 2 spliceosome | - | - | 5.93E-04 | 13 |
| proton-transporting ATP synthase complex | - | - | 1.06E-03 | 11 |
| mitochondrial proton-transporting ATP synthase complex | - | - | 2.37E-03 | 10 |
| DNA replication factor C complex | - | - | 3.93E-03 | 5 |
| U4/U6 x U5 tri-snRNP complex | - | - | 4.34E-03 | 11 |
| spliceosomal tri-snRNP complex | - | - | 4.34E-03 | 11 |
| tricarboxylic acid cycle enzyme complex | - | - | 4.34E-03 | 9 |

| | | | | |
|---|---|---|---|---|
| protein folding chaperone complex | - | - | 4.48E-03 | 13 |
| proton-transporting ATP synthase complex, coupling factor F(o) | - | - | 5.85E-03 | 8 |
| exoribonuclease complex | - | - | 6.51E-03 | 9 |
| nucleocytoplasmic transport complex | - | - | 7.66E-03 | 7 |
| proton-transporting two-sector ATPase complex, proton-transporting domain | - | - | 1.03E-02 | 10 |
| MCM complex | - | - | 1.03E-02 | 7 |
| mitochondrial proton-transporting ATP synthase complex, coupling factor F(o) | - | - | 1.04E-02 | 5 |
| nucleolar exosome (RNase complex) | - | - | 1.14E-02 | 5 |
| smooth endoplasmic reticulum | - | - | 1.27E-02 | 12 |
| exosome (RNase complex) | - | - | 1.29E-02 | 8 |
| DNA replication preinitiation complex | - | - | 1.57E-02 | 7 |
| mitochondrial tricarboxylic acid cycle enzyme complex | - | - | 1.64E-02 | 6 |
| laminin-10 complex | - | - | 2.08E-02 | 3 |
| signal peptidase complex | - | - | 2.09E-02 | 4 |

| | | | | |
|---|---|---|---|---|
| multi-eIF complex | - | - | 2.41E-02 | 3 |
| cytoplasmic exosome (RNase complex) | - | - | 2.49E-02 | 6 |
| intercellular canaliculus | - | - | 2.52E-02 | 4 |
| CMG complex | - | - | 2.57E-02 | 6 |
| NLS-dependent protein nuclear import complex | - | - | 2.65E-02 | 5 |
| nuclear exosome (RNase complex) | - | - | 2.66E-02 | 6 |
| eukaryotic translation elongation factor 1 complex | - | - | 2.66E-02 | 4 |
| proteasome complex | - | - | 2.78E-02 | 16 |
| perichromatin fibrils | - | - | 2.81E-02 | 4 |
| proton-transporting ATP synthase complex, catalytic core F(1) | - | - | 2.81E-02 | 4 |
| RNA polymerase II, core complex | - | - | 3.46E-02 | 6 |
| DNA replication factor A complex | - | - | 3.66E-02 | 5 |
| endocytic vesicle lumen | - | - | 3.81E-02 | 9 |
| protein kinase CK2 complex | - | - | 3.85E-02 | 3 |
| mRNA cleavage factor complex | - | - | 3.85E-02 | 9 |
| mitochondrial outer membrane translocase complex | - | - | 4.17E-02 | 7 |

| | | | | |
|---|---|---|---|---|
| organellar ribosome | - | - | 4.99E-02 | 15 |
| mitochondrial ribosome | - | - | 4.99E-02 | 15 |

**FDR**: False discovery rate

**WT:** Wild-type

**Supplementary Table S3. Significant pathways resulted from pathway analysis of significantly differential metabolites between isoniazid-treated and untreated wild-type HepG2 cells.**

| KEGG Pathway | Pathway Size | WT – 17 mM vs. WT – 0 mM | | WT – 32.5 mM vs. WT – 0 mM | |
|---|---|---|---|---|---|
| | | Significant Hits | P-value | Significant Hits | P-value |
| Purine metabolism | 70 | 3 | 2.15E-02 | - | - |
| Glutathione metabolism | 28 | 2 | 2.43E-02 | - | - |
| Arginine and proline metabolism | 36 | 2 | 3.89E-02 | - | - |
| Glyoxylate and dicarboxylate metabolism | 32 | - | - | 4 | 1.32E-03 |
| Nicotinate and nicotinamide metabolism | 15 | - | - | 3 | 1.42E-03 |
| Pyrimidine metabolism | 39 | - | - | 4 | 2.80E-03 |
| Citrate cycle (TCA cycle) | 20 | - | - | 3 | 3.37E-03 |
| Alanine, aspartate and glutamate metabolism | 28 | - | - | 3 | 8.91E-03 |
| Arginine biosynthesis | 14 | - | - | 2 | 1.96E-02 |
| Histidine metabolism | 16 | - | - | 2 | 2.53E-02 |
| Pantothenate and CoA biosynthesis | 20 | - | - | 2 | 3.86E-02 |
| beta-Alanine metabolism | 21 | - | - | 2 | 4.23E-02 |

**KEGG:** Kyoto Encyclopedia of Genes and Genomes

**WT:** Wild-type

**Supplementary Table S4. Significant pathways resulted from over-representation analysis of significantly differential proteins between Nrf2-knockout and wild-type HepG2 cells upon isoniazid treatments.**

| KEGG Pathway | FDR | Number of Enriched Proteins |
|---|---|---|
| Carbon metabolism | 4.11E-14 | 49 |
| Protein processing in endoplasmic reticulum | 4.61E-14 | 61 |
| Biosynthesis of amino acids | 8.71E-12 | 35 |
| Glycolysis / Gluconeogenesis | 4.14E-08 | 28 |
| Spliceosome | 4.14E-08 | 59 |
| Valine, leucine and isoleucine degradation | 4.35E-08 | 23 |
| Fatty acid degradation | 1.32E-07 | 21 |
| Biosynthesis of cofactors | 1.76E-07 | 45 |
| Glutathione metabolism | 4.82E-07 | 24 |
| Pyruvate metabolism | 4.06E-06 | 20 |
| Peroxisome | 4.75E-06 | 28 |
| Fatty acid metabolism | 6.41E-06 | 22 |
| Arginine and proline metabolism | 1.06E-05 | 20 |
| Cysteine and methionine metabolism | 2.07E-05 | 20 |
| Nucleocytoplasmic transport | 4.31E-05 | 31 |
| Bacterial invasion of epithelial cells | 4.41E-05 | 25 |
| Citrate cycle (TCA cycle) | 5.00E-05 | 14 |
| Nucleotide metabolism | 6.59E-05 | 26 |
| Chemical carcinogenesis - reactive oxygen species | 7.31E-05 | 51 |
| Parkinson disease | 1.90E-04 | 57 |
| Huntington disease | 2.24E-04 | 63 |
| Salmonella infection | 3.12E-04 | 53 |
| Pentose phosphate pathway | 3.40E-04 | 13 |

| | | |
|---|---|---|
| Protein export | 3.40E-04 | 13 |
| Amyotrophic lateral sclerosis | 4.08E-04 | 71 |
| Propanoate metabolism | 4.71E-04 | 13 |
| Amino sugar and nucleotide sugar metabolism | 4.71E-04 | 17 |
| Biosynthesis of nucleotide sugars | 5.57E-04 | 14 |
| Drug metabolism - other enzymes | 5.57E-04 | 23 |
| Lysosome | 6.63E-04 | 32 |
| Fructose and mannose metabolism | 8.40E-04 | 13 |
| Ascorbate and aldarate metabolism | 9.05E-04 | 12 |
| Pyrimidine metabolism | 9.05E-04 | 18 |
| Pathogenic Escherichia coli infection | 1.11E-03 | 43 |
| Diabetic cardiomyopathy | 1.36E-03 | 43 |
| Tyrosine metabolism | 1.43E-03 | 13 |
| Alanine, aspartate and glutamate metabolism | 1.91E-03 | 13 |
| Tryptophan metabolism | 1.98E-03 | 14 |
| Ferroptosis | 1.98E-03 | 14 |
| 2-Oxocarboxylic acid metabolism | 2.13E-03 | 12 |
| Sulfur metabolism | 3.18E-03 | 6 |
| Histidine metabolism | 4.36E-03 | 9 |
| Fatty acid biosynthesis | 4.36E-03 | 8 |
| beta-Alanine metabolism | 4.36E-03 | 11 |
| Glyoxylate and dicarboxylate metabolism | 4.36E-03 | 11 |
| Pentose and glucuronate interconversions | 4.65E-03 | 12 |
| Glycine, serine and threonine metabolism | 4.72E-03 | 13 |
| Pathways of neurodegeneration - multiple diseases | 5.84E-03 | 81 |
| Prion disease | 7.35E-03 | 51 |
| Vibrio cholerae infection | 1.35E-02 | 14 |

| | | |
|---|---|---|
| Alzheimer disease | 1.38E-02 | 66 |
| Platinum drug resistance | 1.66E-02 | 18 |
| Thyroid hormone synthesis | 1.66E-02 | 18 |
| Shigellosis | 1.73E-02 | 45 |
| Fatty acid elongation | 1.83E-02 | 9 |
| Central carbon metabolism in cancer | 2.09E-02 | 17 |
| Arginine biosynthesis | 2.18E-02 | 8 |
| Renin-angiotensin system | 2.18E-02 | 8 |
| Aminoacyl-tRNA biosynthesis | 2.26E-02 | 16 |
| HIF-1 signaling pathway | 2.40E-02 | 23 |
| Purine metabolism | 2.40E-02 | 26 |
| Metabolism of xenobiotics by cytochrome P450 | 2.57E-02 | 18 |
| Legionellosis | 2.70E-02 | 14 |
| Apoptosis | 2.70E-02 | 27 |
| Oxidative phosphorylation | 3.25E-02 | 27 |
| One carbon pool by folate | 3.25E-02 | 7 |
| PPAR signaling pathway | 3.63E-02 | 17 |
| Phenylalanine metabolism | 3.88E-02 | 6 |
| Pantothenate and CoA biosynthesis | 4.20E-02 | 7 |
| Regulation of actin cytoskeleton | 4.29E-02 | 40 |
| Selenocompound metabolism | 4.95E-02 | 6 |
| Drug metabolism - cytochrome P450 | 4.95E-02 | 16 |
| Butanoate metabolism | 4.95E-02 | 8 |
| Folate biosynthesis | 4.95E-02 | 8 |
| Biosynthesis of unsaturated fatty acids | 4.95E-02 | 8 |

**KEGG:** Kyoto Encyclopedia of Genes and Genomes

**FDR**: False discovery rate

**Supplementary Table S5. Significant gene ontology (cellular component) terms resulted from over-representation analysis of significantly differential proteins between Nrf2-knockout and wild-type HepG2 cells upon isoniazid treatments.**

| Terms | FDR | Number of Enriched Proteins |
|---|---|---|
| mitochondrial matrix | 1.04E-30 | 132 |
| focal adhesion | 7.75E-27 | 114 |
| cell-substrate junction | 4.96E-26 | 114 |
| cytoplasmic vesicle lumen | 6.48E-22 | 90 |
| vesicle lumen | 6.57E-22 | 90 |
| secretory granule lumen | 9.00E-22 | 89 |
| preribosome | 2.74E-18 | 45 |
| ficolin-1-rich granule | 2.74E-18 | 60 |
| spliceosomal complex | 1.47E-17 | 63 |
| ficolin-1-rich granule lumen | 8.41E-16 | 45 |
| catalytic step 2 spliceosome | 1.01E-15 | 38 |
| endoplasmic reticulum lumen | 3.03E-15 | 76 |
| melanosome | 7.24E-14 | 40 |
| pigment granule | 7.24E-14 | 40 |
| primary lysosome | 3.06E-13 | 47 |
| azurophil granule | 3.06E-13 | 47 |
| endoplasmic reticulum protein-containing complex | 1.05E-12 | 41 |
| mitochondrial inner membrane | 1.68E-12 | 96 |
| U2-type spliceosomal complex | 6.60E-12 | 34 |
| vacuolar lumen | 1.09E-11 | 48 |
| coated vesicle | 1.49E-11 | 69 |
| small-subunit processome | 2.35E-11 | 29 |

| | | |
|---|---|---|
| endoplasmic reticulum-Golgi intermediate compartment | 2.90E-11 | 40 |
| coated vesicle membrane | 7.92E-11 | 51 |
| organelle outer membrane | 1.35E-10 | 57 |
| outer membrane | 1.83E-10 | 57 |
| vesicle coat | 5.26E-10 | 25 |
| mitochondrial protein-containing complex | 1.19E-09 | 63 |
| peroxisome | 1.19E-09 | 39 |
| microbody | 1.19E-09 | 39 |
| endoplasmic reticulum chaperone complex | 5.08E-09 | 10 |
| preribosome, large subunit precursor | 5.08E-09 | 13 |
| azurophil granule lumen | 5.70E-09 | 29 |
| cortical cytoskeleton | 8.91E-09 | 31 |
| cortical actin cytoskeleton | 1.18E-08 | 25 |
| nuclear envelope | 1.47E-08 | 85 |
| nuclear periphery | 1.75E-08 | 38 |
| mitochondrial outer membrane | 2.33E-08 | 48 |
| nuclear speck | 2.70E-08 | 74 |
| ribonucleoprotein granule | 9.09E-08 | 54 |
| cytoplasmic stress granule | 1.42E-07 | 27 |
| U2-type precatalytic spliceosome | 2.04E-07 | 19 |
| Sm-like protein family complex | 2.06E-07 | 31 |
| small nuclear ribonucleoprotein complex | 2.19E-07 | 29 |
| organelle envelope lumen | 3.45E-07 | 26 |
| spliceosomal snRNP complex | 3.45E-07 | 27 |
| nuclear matrix | 4.03E-07 | 32 |
| membrane coat | 4.05E-07 | 27 |
| coated membrane | 4.05E-07 | 27 |

| | | |
|---|---|---|
| cytoplasmic ribonucleoprotein granule | 4.05E-07 | 50 |
| cell cortex | 4.25E-07 | 57 |
| COPI-coated vesicle membrane | 4.56E-07 | 11 |
| mitochondrial intermembrane space | 4.56E-07 | 24 |
| precatalytic spliceosome | 4.84E-07 | 19 |
| tertiary granule | 5.54E-07 | 37 |
| U2-type catalytic step 2 spliceosome | 5.80E-07 | 14 |
| COPI vesicle coat | 1.65E-06 | 9 |
| lysosomal lumen | 1.80E-06 | 26 |
| U2 snRNP | 1.81E-06 | 13 |
| cell leading edge | 2.80E-06 | 69 |
| 90S preribosome | 2.92E-06 | 13 |
| COPI-coated vesicle | 3.31E-06 | 12 |
| fibrillar center | 4.74E-06 | 33 |
| oxidoreductase complex | 6.24E-06 | 30 |
| lamellipodium | 6.43E-06 | 40 |
| Golgi-associated vesicle membrane | 8.01E-06 | 18 |
| Golgi-associated vesicle | 9.96E-06 | 24 |
| nuclear membrane | 1.55E-05 | 53 |
| COPII vesicle coat | 1.72E-05 | 9 |
| tertiary granule lumen | 2.13E-05 | 17 |
| chromosomal region | 2.45E-05 | 63 |
| methylosome | 2.55E-05 | 8 |
| U5 snRNP | 2.61E-05 | 11 |
| U4/U6 x U5 tri-snRNP complex | 2.77E-05 | 15 |
| peroxisomal matrix | 3.14E-05 | 16 |
| microbody lumen | 3.14E-05 | 16 |

| | | |
|---|---|---|
| spliceosomal tri-snRNP complex | 3.64E-05 | 15 |
| COP9 signalosome | 4.28E-05 | 13 |
| peroxisomal membrane | 4.28E-05 | 18 |
| microbody membrane | 4.28E-05 | 18 |
| ribosomal subunit | 6.86E-05 | 36 |
| clathrin-coated vesicle | 8.12E-05 | 39 |
| ribosome | 8.90E-05 | 43 |
| COPII-coated ER to Golgi transport vesicle | 1.02E-04 | 22 |
| inner mitochondrial membrane protein complex | 1.61E-04 | 31 |
| transport vesicle | 2.41E-04 | 63 |
| midbody | 2.51E-04 | 36 |
| blood microparticle | 2.85E-04 | 28 |
| specific granule | 3.11E-04 | 30 |
| endoplasmic reticulum exit site | 3.48E-04 | 11 |
| trans-Golgi network transport vesicle | 4.06E-04 | 12 |
| lysosomal membrane | 4.15E-04 | 63 |
| lytic vacuole membrane | 4.15E-04 | 63 |
| large ribosomal subunit | 4.46E-04 | 24 |
| intercalated disc | 5.38E-04 | 14 |
| respirasome | 5.49E-04 | 22 |
| ER to Golgi transport vesicle membrane | 5.71E-04 | 16 |
| chromosome, centromeric region | 6.42E-04 | 41 |
| SMN-Sm protein complex | 6.75E-04 | 8 |
| U12-type spliceosomal complex | 6.75E-04 | 10 |
| peptidase complex | 7.22E-04 | 24 |
| endopeptidase complex | 7.44E-04 | 19 |
| clathrin-coated vesicle membrane | 7.88E-04 | 26 |

| | | |
|---|---|---|
| cell division site | 8.28E-04 | 17 |
| cell-cell contact zone | 8.28E-04 | 17 |
| platelet alpha granule lumen | 9.38E-04 | 16 |
| vacuolar membrane | 1.04E-03 | 66 |
| ficolin-1-rich granule membrane | 1.04E-03 | 15 |
| secretory granule membrane | 1.13E-03 | 47 |
| lipid droplet | 1.20E-03 | 21 |
| proteasome complex | 1.23E-03 | 15 |
| U2-type catalytic step 1 spliceosome | 1.48E-03 | 6 |
| catalytic step 1 spliceosome | 1.48E-03 | 6 |
| filopodium | 1.52E-03 | 21 |
| methyltransferase complex | 1.84E-03 | 20 |
| azurophil granule membrane | 1.95E-03 | 14 |
| respiratory chain complex | 2.20E-03 | 19 |
| sno(s)RNA-containing ribonucleoprotein complex | 2.33E-03 | 9 |
| chromosome, telomeric region | 2.33E-03 | 29 |
| U2-type prespliceosome | 2.76E-03 | 7 |
| prespliceosome | 2.76E-03 | 7 |
| membrane microdomain | 2.87E-03 | 42 |
| organellar ribosome | 2.91E-03 | 18 |
| mitochondrial ribosome | 2.91E-03 | 18 |
| nuclear pore | 3.09E-03 | 19 |
| mitochondrial respirasome | 3.09E-03 | 19 |
| Prp19 complex | 3.63E-03 | 6 |
| cytolytic granule | 3.63E-03 | 6 |
| specific granule lumen | 3.63E-03 | 14 |
| actomyosin | 3.67E-03 | 18 |

| | | |
|---|---|---|
| smooth endoplasmic reticulum | 3.73E-03 | 9 |
| brush border | 4.01E-03 | 20 |
| organellar large ribosomal subunit | 4.05E-03 | 13 |
| mitochondrial large ribosomal subunit | 4.05E-03 | 13 |
| rough endoplasmic reticulum | 4.05E-03 | 13 |
| P-body | 4.23E-03 | 19 |
| contractile ring | 4.23E-03 | 5 |
| membrane raft | 4.44E-03 | 41 |
| spindle | 4.44E-03 | 57 |
| nuclear outer membrane | 4.54E-03 | 9 |
| ruffle | 4.58E-03 | 29 |
| ATPase complex | 4.69E-03 | 25 |
| mitochondrial respiratory chain complex I | 5.23E-03 | 12 |
| NADH dehydrogenase complex | 5.23E-03 | 12 |
| respiratory chain complex I | 5.23E-03 | 12 |
| sarcoplasm | 5.53E-03 | 18 |
| transport vesicle membrane | 6.13E-03 | 35 |
| endoplasmic reticulum-Golgi intermediate compartment membrane | 6.36E-03 | 16 |
| proteasome core complex, beta-subunit complex | 6.62E-03 | 5 |
| U1 snRNP | 6.91E-03 | 9 |
| nucleoid | 6.91E-03 | 11 |
| mitochondrial nucleoid | 6.91E-03 | 11 |
| endoplasmic reticulum quality control compartment | 6.98E-03 | 6 |
| Cajal body | 6.98E-03 | 16 |
| adherens junction | 6.98E-03 | 29 |
| condensed chromosome, centromeric region | 7.93E-03 | 28 |
| platelet alpha granule | 8.14E-03 | 17 |

| | | |
|---|---|---|
| cleavage furrow | 8.14E-03 | 12 |
| site of DNA damage | 8.22E-03 | 21 |
| telomerase holoenzyme complex | 8.22E-03 | 7 |
| sarcoplasmic reticulum | 8.23E-03 | 15 |
| condensed chromosome | 9.10E-03 | 39 |
| sarcolemma | 9.32E-03 | 23 |
| eukaryotic 48S preinitiation complex | 9.32E-03 | 6 |
| RNA cap binding complex | 9.32E-03 | 6 |
| cytoplasmic ubiquitin ligase complex | 9.35E-03 | 5 |
| transcription export complex | 9.35E-03 | 5 |
| mitotic spindle | 9.35E-03 | 28 |
| stress fiber | 9.35E-03 | 16 |
| contractile actin filament bundle | 9.35E-03 | 16 |
| endocytic vesicle | 1.11E-02 | 46 |
| actin filament bundle | 1.20E-02 | 17 |
| kinetochore | 1.20E-02 | 26 |
| glutamatergic synapse | 1.22E-02 | 52 |
| SAM complex | 1.33E-02 | 5 |
| U4 snRNP | 1.33E-02 | 5 |
| eukaryotic translation initiation factor 4F complex | 1.33E-02 | 5 |
| transcription preinitiation complex | 1.33E-02 | 5 |
| MIB complex | 1.33E-02 | 5 |
| plasma membrane raft | 1.39E-02 | 19 |
| organelle membrane contact site | 1.39E-02 | 11 |
| translation preinitiation complex | 1.60E-02 | 6 |
| proton-transporting two-sector ATPase complex | 1.61E-02 | 11 |
| uropod | 1.85E-02 | 5 |

| | | |
|---|---|---|
| septin ring | 1.85E-02 | 5 |
| septin complex | 1.85E-02 | 5 |
| phagocytic vesicle | 1.91E-02 | 22 |
| endocytic vesicle membrane | 1.94E-02 | 29 |
| proteasome core complex | 2.02E-02 | 6 |
| trans-Golgi network transport vesicle membrane | 2.02E-02 | 6 |
| endolysosome membrane | 2.02E-02 | 6 |
| clathrin-coated endocytic vesicle | 2.09E-02 | 16 |
| podosome | 2.15E-02 | 8 |
| actin filament | 2.30E-02 | 19 |
| dendrite cytoplasm | 2.40E-02 | 7 |
| cell trailing edge | 2.44E-02 | 5 |
| retromer complex | 2.54E-02 | 6 |
| small ribosomal subunit | 2.55E-02 | 14 |
| microtubule | 2.72E-02 | 56 |
| cytosolic ribosome | 2.92E-02 | 19 |
| endoribonuclease complex | 3.00E-02 | 8 |
| phagocytic vesicle membrane | 3.12E-02 | 14 |
| mitochondrial outer membrane translocase complex | 3.13E-02 | 6 |
| mitochondria-associated endoplasmic reticulum membrane | 3.13E-02 | 6 |
| septin cytoskeleton | 3.16E-02 | 5 |
| caveola | 3.41E-02 | 14 |
| endolysosome | 3.41E-02 | 7 |
| site of polarized growth | 3.53E-02 | 24 |
| aminoacyl-tRNA synthetase multienzyme complex | 3.71E-02 | 4 |
| mitochondrial tricarboxylic acid cycle enzyme complex | 3.71E-02 | 4 |
| clathrin-coated endocytic vesicle membrane | 3.75E-02 | 13 |

| | | |
|---|---|---|
| endocytic vesicle lumen | 3.75E-02 | 6 |
| outer mitochondrial membrane protein complex | 3.75E-02 | 6 |
| ruffle membrane | 3.75E-02 | 16 |
| actin-based cell projection | 3.85E-02 | 30 |
| immunological synapse | 4.33E-02 | 9 |
| late endosome | 4.33E-02 | 39 |
| growth cone | 4.46E-02 | 23 |
| clathrin-coated pit | 4.48E-02 | 13 |
| rough endoplasmic reticulum membrane | 4.51E-02 | 6 |
| nuclear lamina | 4.87E-02 | 4 |
| eukaryotic 43S preinitiation complex | 4.87E-02 | 5 |
| clathrin coat of trans-Golgi network vesicle | 4.87E-02 | 5 |
| clathrin coat of coated pit | 4.87E-02 | 5 |
| pseudopodium | 4.87E-02 | 5 |
| tricarboxylic acid cycle enzyme complex | 4.87E-02 | 5 |

**FDR**: False discovery rate

**Supplementary Table S6. Significant pathways resulted from pathway analysis of significantly differential metabolites between Nrf2-knockout and wild-type HepG2 cells upon isoniazid treatments.**

| KEGG Pathways | Pathway Size | Significant Hits | P-Value |
|---|---|---|---|
| Alanine, aspartate and glutamate metabolism | 28 | 2 | 7.99E-03 |
| Glutathione metabolism | 28 | 2 | 7.99E-03 |
| Purine metabolism | 70 | 2 | 4.58E-02 |

**KEGG:** Kyoto Encyclopedia of Genes and Genomes

**Reference**


1. Park S.J., Lee S.B., Suh Y., Kim S.J., Lee N., Hong J.H., et al. DISC1 Modulates Neuronal Stress Responses by Gate-Keeping ER-Mitochondria Ca(2+) Transfer through the MAM. Cell Rep. 2017;21(10):2748-59.
2. Imamura H., Nhat K.P., Togawa H., Saito K., Iino R., Kato-Yamada Y., et al. Visualization of ATP levels inside single living cells with fluorescence resonance energy transfer-based genetically encoded indicators. Proc Natl Acad Sci U S A. 2009;106(37):15651-6.
3. Ermakova Y.G., Bilan D.S., Matlashov M.E., Mishina N.M., Markvicheva K.N., Subach O.M., et al. Red fluorescent genetically encoded indicator for intracellular hydrogen peroxide. Nat Commun. 2014;5:5222.
4. Gutscher M., Pauleau A.L., Marty L., Brach T., Wabnitz G.H., Samstag Y., et al. Real-time imaging of the intracellular glutathione redox potential. Nat Methods. 2008;5(6):553-9.
5. Dagda R.K., Cherra S.J., 3rd, Kulich S.M., Tandon A., Park D., and Chu C.T. Loss of PINK1 function promotes mitophagy through effects on oxidative stress and mitochondrial fission. J Biol Chem. 2009;284(20):13843-55.
6. Cajka T., Hricko J., Rudl Kulhava L., Paucova M., Novakova M., and Kuda O. Optimization of Mobile Phase Modifiers for Fast LC-MS-Based Untargeted Metabolomics and Lipidomics. Int J Mol Sci. 2023;24(3).
7. Yen N.T.H., Anh N.K., Jayanti R.P., Phat N.K., Vu D.H., Ghim J.L., et al. Multimodal plasma metabolomics and lipidomics in elucidating metabolic perturbations in tuberculosis patients with concurrent type 2 diabetes. Biochimie. 2023;211:153-63.
8. Anh N.K., Yen N.T.H., Tien N.T.N., Phat N.K., Park Y.J., Kim H.S., et al. Metabolic phenotyping and global functional analysis facilitate metabolic signature discovery for tuberculosis treatment monitoring. Biochim Biophys Acta Mol Basis Dis. 2024;1870(4):167064.
9. Anh N.K., Phat N.K., Thu N.Q., Tien N.T.N., Eunsu C., Kim H.S., et al. Discovery of urinary biosignatures for tuberculosis and nontuberculous mycobacteria classification using metabolomics and machine learning. Sci Rep. 2024;14(1):15312.
10. Thi My Nhung T., Phuoc Long N., Diem Nghi T., Suh Y., Hoang Anh N., Jung C.W., et al. Genome-wide kinase-MAM interactome screening reveals the role of CK2A1 in MAM Ca(2+) dynamics linked to DEE66. Proc Natl Acad Sci U S A. 2023;120(32):e2303402120.
11. Nguyen B.T., Le Q.V., Ahn J., Nguyen K.A., Nguyen H.T., Kang J.S., et al. Omics analysis unveils changes in the metabolome and lipidome of Caenorhabditis elegans upon polydopamine exposure. J Pharm Biomed Anal. 2024;244:116126.
12. Lavarello C., Barco S., Bartolucci M., Panfoli I., Magi E., Tripodi G., et al. Development of an Accurate Mass Retention Time Database for Untargeted Metabolomic Analysis and Its Application to Plasma and Urine Pediatric Samples. Molecules. 2021;26(14).
13. Tsugawa H., Ikeda K., Takahashi M., Satoh A., Mori Y., Uchino H., et al. A lipidome atlas in MS-DIAL 4. Nat Biotechnol. 2020;38(10):1159-63.

14. Kind T., Liu K.H., Lee D.Y., DeFelice B., Meissen J.K., and Fiehn O. LipidBlast in silico tandem mass spectrometry database for lipid identification. Nat Methods. 2013;10(8):755-8.
15. Hutchings C., Dawson C.S., Krueger T., Lilley K.S., and Breckels L.M. A Bioconductor workflow for processing, evaluating, and interpreting expression proteomics data. F1000Res. 2023;12:1402.
16. Pang Z., Lu Y., Zhou G., Hui F., Xu L., Viau C., et al. MetaboAnalyst 6.0: towards a unified platform for metabolomics data processing, analysis and interpretation. Nucleic Acids Res. 2024;52(W1):W398-W406.
17. Gu Z. Complex heatmap visualization. Imeta. 2022;1(3):e43.
18. Yu G., Wang L.G., Han Y., and He Q.Y. clusterProfiler: an R package for comparing biological themes among gene clusters. OMICS. 2012;16(5):284-7.
19. Kanehisa M. and Goto S. KEGG: kyoto encyclopedia of genes and genomes. Nucleic Acids Res. 2000;28(1):27-30.
20. Molenaar M.R., Jeucken A., Wassenaar T.A., van de Lest C.H.A., Brouwers J.F., and Helms J.B. LION/web: a web-based ontology enrichment tool for lipidomic data analysis. Gigascience. 2019;8(6).
